# Bacterial plasmid-associated and chromosomal proteins have fundamentally different properties in protein interaction networks

**DOI:** 10.1101/2022.04.29.490008

**Authors:** Tim Downing, Alexander Rahm

## Abstract

Plasmids facilitate horizontal gene transfer, which enables the diversification of pathogens into new anatomical and environmental niches, implying that plasmid-encoded genes can cooperate well with chromosomal genes. We hypothesise that such mobile genes are functionally different to chromosomal ones due to this ability to encode non-essential functions like antimicrobial resistance and traverse distinct host cells. The effect of plasmid-driven gene gain on protein-protein interaction network topology is an important question in this area. Moreover, the extent to which these chromosomally- and plasmid-encoded proteins interact with proteins from their own groups compared to the levels with the other group remains unclear. Here, we examined the incidence and protein-protein interactions of all known plasmid-encoded genes across representative specimens from most bacteria using all available plasmids. We found that such plasmid-encoded genes constitute ∼0.7% of the total number of genes per bacterial sample, and that plasmid genes are preferentially associated with different species but had limited taxonomical power beyond this. Surprisingly, plasmid-encoded proteins had both more protein-protein interactions compared to chromosomal proteins, countering the hypothesis that genes with higher mobility rates should have fewer protein-level interactions. Nonetheless, topological analysis and investigation of the protein-protein interaction networks’ connectivity and change in the number of independent components demonstrated that the plasmid-encoded proteins had limited overall impact in >96% of samples. This paper assembled extensive data on plasmid-encoded proteins, their interactions and associations with diverse bacterial specimens that is available for the community to investigate in more detail.

**Significance statement:** It is well-established that plasmids drive new traits in their bacterial hosts, but the extent to which host-plasmid co-evolution is evident at the level of protein-protein interactions remains unclear. To address this, we compiled and analysed all available valid bacterial plasmids and associated proteins to explore the compositional differences between chromosomal and plasmid-encoded proteins and their interaction levels. We found that plasmid-encoded genes were highly correlated across the bacterial samples such that they had a high association with taxonomic context. Contrasting with the complexity hypothesis, plasmid-encoded proteins had far more interactions on average than chromosomal ones, though they had minimal effects on protein-protein interaction network structure. This demonstrated that host-plasmid co-evolution is evident and detectable at the level of protein interactions.

## Introduction

Plasmids are short extrachromosomal DNA elements that are typically circular and with variable copy numbers. Conjugative plasmids possess machinery enabling horizontal gene transfer (HGT) between bacterial cells, a key mechanism of bacterial evolution. Plasmid DNA may also transfer infrequently between cells by transformation, vesicles and phages (Canosi et al 1982, Erdmann et al 2017, Zhang et al 2018, Wein & Dagan 2020). This allows plasmid-encoded genes to move between bacterial cells in the same niche (Wein et al 2021), where these genes may allow new phenotypes (Norman et al 2009, Downing 2015, Hall et al 2017, Decano et al 2020). Plasmid-encoded genes encode a wide range of functions, including antimicrobial resistance (AMR), virulence, metabolism, and symbiosis (Ahmer et al 1999, Stasiak et al 2014, San Millan 2018), facilitating spread into new environmental niches (Niehus et al 2015). Importantly, they depend on the environmental context to be beneficial, such that they can be lost when no longer required (Niehus et al 2015).

Mobile genes form part of the accessory genome, distinct from chromosomal genes that typically compose the core genome (Brockhurst et al 2019), though some chromosomal genes may not be universal within a collection, and so are part of the accessory genome. Some genes can be encoded on both plasmids and chromosomes, and there are numerous instances in which beneficial genes have been mobilised to spread from plasmids to chromosomes, such as *bla*CTX-M-15 (Agyekum et al 2016, Huang et al 2017, Irrgang et al 2017, Decano et al 2019a, Decano et al 2019b, Yoon et al 2020, Ludden et al 2020, Bevan et al 2021, Shawa et al 2021). In this study, we examined the broad patterns across bacteria using the categorisation of plasmid-encoded versus chromosomal.

Plasmids impose a metabolic fitness cost on their host cells (San Millan & MacLean 2017, Baltrus 2013, Harrison & Brockhurst 2012) and yet persist widely thanks to infectious spread, stability mechanisms, and chromosomal compensatory mutations (Stalder et al 2017). Plasmids are genetically very diverse, and this is paralleled by extensive variation in AMR gene profiles across bacteria (Ho et al 2020, Coelho et al 2021). The co-evolution of plasmid-encoded genes with chromosomal ones are consequently of keen interest. This matters because a suitable host-recipient genetic background is needed such that the transferred gene can be acquired and replicated (Forsberg et al 2014, Soucy et al 2015, Andam & Gogarten 2011). In addition, donor & recipient cells may be genetically distinct and yet have high HGT rates (Smillie et al 2011, Brito et al 2016), suggesting additional factors affecting gene and plasmid retention to quantify.

HGT depends on the physiological compatibility so that a plasmid-bourne gene can be expressed and useful in the transconjugant recipient cells by interacting with host proteins in a beneficial manner (Porse et al 2018). The complexity hypothesis (Jain et al 1999, Novick & Doolittle 2020) asserts that protein-protein interaction (PPI) network connectivity and HGT rates are negatively correlated. In line with this, previous work on small sets of protein orthologs has found that less essential genes with higher HGT rates have fewer PPIs (Nakamura et al 2004, Aris-Brosou 2005, Puigbo et al 2010). HGT rates are higher for rarer genes like those in the accessory genome on plasmids, and lower for core genes (Dewar et al 2021, Touchon et al 2020). Moreover, gene context and physical gene distance are related to network proximity (Babu et al 2006) such that accessory genes and newly transferred genes are likely to be peripheral in PPI networks, including plasmid-encoded ones, whereas the chromosomal proteins should be more central.

Plasmid properties vary extensively based on the host cell (De Gelder et al 2007, Dunn et al 2021, Alonso-Del Valle et al 2021, Kottara et al 2018, Sheppard et al 2020, Gama et al 2020, Alderliesten et al 2020), suggesting that the interaction between host chromosomal proteins with plasmid-encoded ones is important. The evidence collated so far implies that chromosomes adapt to plasmids more than the reverse, typically by single nucleotide polymorphisms (SNPs) (Stalder et al 2017, Harrison et al 2015, San Millan et al 2014, Hall et al 2020), but sometimes by larger gene deletions (Modi et al 1991, Porse et al 2016, Lee & Marx 2012). This co-evolution of plasmid-encoded and chromosomal proteins means more systematic approaches to examining the extent to which these proteins interact across bacterial species could provide insights into plasmid-host compatibility.

In this study, we assessed the hypothesis that chromosomal and plasmid-encoded genes have different interaction levels based on the host species, PPI context and PPI connectivity. Using a core dataset of 4,363 bacterial samples, we examined PPI network connectivity using PPI data for all chromosomal proteins and for plasmid-encoded ones. Importantly, HGT is not strongly affected by gene sequence composition and features (Porse et al 2018), supporting our simple classification of genes as present or absent here. We examined the changes in PPI network structure based on changes in indirect connections when plasmid-encoded proteins enter a host’s chromosomal PPI network. We identified distinctive features associated with plasmid-encoded proteins and their PPIs, assessed their covariation, and thus provide a pilot study of plasmid-related PPIs s across all bacteria.

## Methods

### Protein-protein interaction data extraction

We retrieved PPI information for all available valid bacterial genomes (n=4,445) from STRING database v12 (Szklarczyk et al 2021) with R v4.0.1 (R Core Team 2021) and RStudio v2022.2.3.492 (RStudio Team 2022) via Bioconductor v3.11 (Ihaka & Gentleman 1996) and packages Rentrez v1.2.2 (Winter 2017), STRINGdb v2.0.1 (Szklarczyk et al 2018), and StringR v1.4.0 (Wickham 2019) (Figure 1). Of the initial 4,445 potential samples listed, 22 were not accessible and four had more >300,000 PPIs, suggesting potential accuracy issues and so they were excluded from analysis. The valid data (n=4,419 samples) was processed and collated using Dplyr v1.0.8 (Wickham et al 2002), Forcats v0.5.1 (Wickham 2021), Grid v4.4.1, ReadR v2.1.2 (Wickham et al 2022), Readxl v1.4.0 (Wickham & Bryan 2022), Tibble v3.1.6 (Müller & Wickham 2021), TidyR v1.2.0 (Wickham & Girlich 2022), Tidyverse v1.3.0 (Wickham et al 2019), VennDiagram v1.7.3 (Chen 2022) and Xlsx v0.6.0. The total number of unique protein names in the entire dataset was 9,551,828, though the true number of gene clusters is likely smaller due to name inaccuracy and redundancy. Using a STRINGdb score threshold of >400, we used this data to compute the numbers of unique proteins and pairwise PPIs on chromosomes: this score threshold of >400 was used throughout this study. These 4,419 post-QC samples had 3,628.5±1,710.0 (mean±SD) proteins and a mean of 57,105.8±34,698.6 PPIs each. Not all data was accessible for 56 samples among these 4,419, leaving 4,363 samples. This and subsequent data below were visualised with R packages ggplot2 v3.3.5 (Wickham 1996) and ggrepel v0.9.1 (Slowikowski 2021). All reported p values were corrected for multiple testing using the Benjamini-Hochberg approach in R.

**Figure 1.**
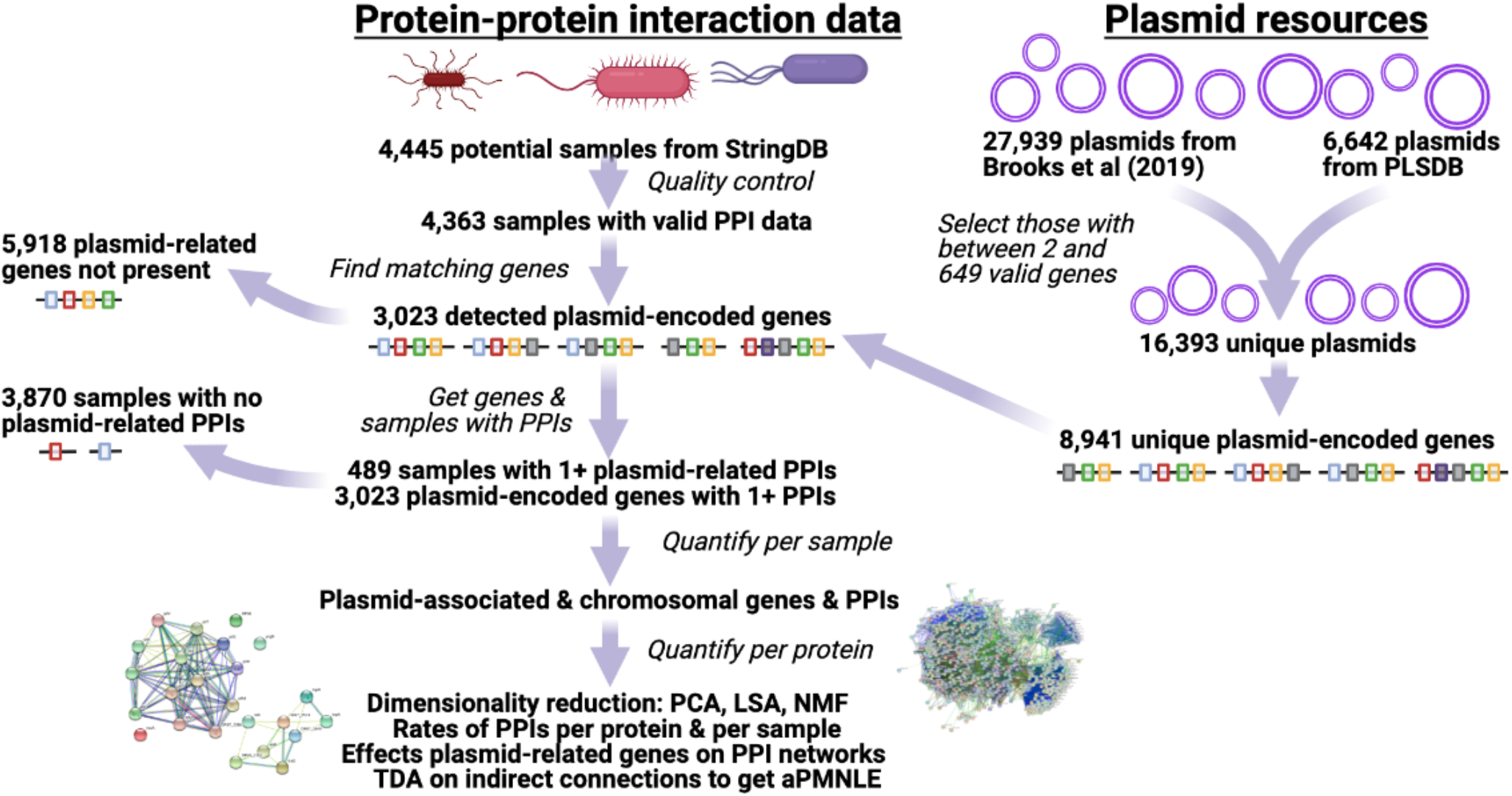
Summary of data sources and analytical steps. The plasmid resources (top right) and protein-protein interaction (PPI) data from StringDB were ultimately merged based on gene matches. Plasmid-encoded genes with PPIs were the focus, and gene prevalence and PPI level per sample were computed, as well as more general patterns across sample based on a range of complementary methods. Created with BioRender.com.

The accuracy of gene names in annotation is a pervasive issue where previous work has shown ∼46±9% of genes can be functionally assigned (Zhou et al 2021). To explore that limiting factor here, we examined the frequency of four-digit (short) gene names to longer ones with five or more digits, after removing any instances of names ending in an underscore or dash or dot followed by any digit. This showed that these samples had a median of 411±410 short names and a median of 2,931±1,744 long names each (Figure S1).

### Identification of plasmid-encoded genes and interactions across bacteria

To get all available plasmid-encoded genes, we collated 32,839 plasmids: 27,939 from Brooks et al (2019) and 6,642 from PLSDB (v2020_11_19, Galata et al 2019) and retrieved gene annotation for each plasmid using Genbankr v1.16.0 (Figure 1). We focused on 16,383 with between two and 640 annotated genes (<640 to avoid chromosome-related contigs). We verified that the plasmids were annotated as plasmids, including 12 with lengths >2 Mb for *Ralstonia solanacearum* and *Rhizobium gallicum*. A gene was defined as plasmid-encoded if it had been found on a plasmid originating in the same species, yielding two categories for genes: plasmid-encoded or chromosomal. These plasmids had a median of 12 unique annotated genes per plasmid and had 8,941 unique plasmid-encoded genes across 98,534 unique sample-gene associations (Table S1, see full table FigShare doi: https://doi.org/10.6084/m9.figshare.19525630). Analyses focused on the 288 samples with at least one plasmid gene, and the 5,538 plasmid-encoded genes detected in at least one sample here.

Next, we identified the numbers of PPIs per protein per sample. A PPI was defined as plasmid-related if either (or both) protein(s) were found on a plasmid, yielding three categories for proteins: chromosomal, plasmid-related with one plasmid-linked protein in the pair, or exclusively plasmid-related where both protein pairs were plasmid-encoded – in practise the latter category was rare and was not investigated in detail here. The numbers of PPIs in which both proteins were exclusively chromosomal was large (286,364,425) compared to the totals for plasmid-related PPIs (390,592) and PPIs exclusively on plasmid-encoded proteins (46,772). This meant the fraction of PPIs that were plasmid-related was 0.136%, and 0.016% of all PPIs were exclusively plasmid-related, and the remainder chromosomal (99.86%). In the samples with at least one plasmid-related PPI, we used the probabilities of having a plasmid-related or chromosomal protein like allele frequencies to compare to the observed frequencies of plasmid-restricted, plasmid-chromosome and chromosome-restricted PPIs per sample in a manner analogous to Wright’s F-statistic (Wright 1951) as *F* = 1 − *obs*/*exp* where *obs* was the fraction of plasmid-chromosome PPIs, and *exp* was twice the product of the frequencies of plasmid-related and chromosomal proteins. This compared the expected probabilities of plasmid-restricted, plasmid-chromosome and chromosome-restricted PPIs against the observed rates to test for structure (or absence thereof) between the plasmid-related and chromosomal proteins.

We examined the taxonomic classifications of the samples across families, orders, classes and phyla using R package taxize v0.9.98 (Chamberlain et al 2020) to examine their scaled pairwise Euclidean distances of the plasmid presence-absence data in a dendrogram from hierarchical clustering from R packages stats v3.6.2 and dendextend v1.15.2 (Galili 2015). This identified totals of 100 families, 46 orders, 20 classes and 10 phyla.

### Screening for correlated bacterial samples and plasmid-encoded genes

We examined patterns of covariance across the set of the samples and 3,023 plasmid-encoded genes using three approaches. To resolve highly correlated sets of samples, the 1^st^ assessed genes and sample-gene pairs using principal components analysis (PCA) implemented with R packages stats v3.6.2 and factoextra v1.0.7 (Kassambara & Mundt 2020). The 2^nd^ was the distributional semantics method, latent semantic indexing (LSA): the probabilistic version has been used to examine other bacteria’s drug resistance genomic data previously (Rusakovica et al 2014). Here, it applied a classical vector space model and singular value decomposition to determine the samples’ plasmid-encoded genes: allocating genes to samples with which they are strongly associated. LSA allocated the samples across the maximum possible 360 dimensions here, which represented the plasmid-encoded genes.

The 3^rd^ approach was non-negative matrix factorisation (NMF) (Lee & Seung 1999) to examine the associations of individual plasmid-encoded genes with individual bacterial samples using R packages nmf v0.23.0 (Gaujoux & Seoighe 2010) and Biobase v2.52.0 (Huber 2015). This was applied to 331 samples with at least three plasmid-encoded genes and 1,733 plasmid-encoded genes found in at three samples as presence-absence data. The optimal rank was determined by examining changes in the cophenetic correlation coefficient across ranks 4-26 for 10 runs per rank to ensure clustering was stable and to select the maximum rank that retained a high cophenetic coefficient. This reduced the complexity of the data per sample to six groups (ranks) (cophenetic correlation coefficient maximised at 0.971). We also examined the covariation of samples with the PPI levels per gene for 481 samples and 2,363 plasmid-encoded genes where the input data was the number of PPIs per protein per sample using NMF as above where each protein had at least 30 PPIs across the sample and each sample had at least ten PPIs in total. The smaller dataset was due to a number of samples and genes with zero plasmid-related PPIs. The optimal rank was determined as above, and an estimate of seven groups (ranks) was obtained (cophenetic correlation coefficient maximised at 0.9856).

### Topological data analysis of the bacterial protein-protein interaction networks

We developed computationally efficient metrics to investigate the topology of a PPI network, with in our focus the PPI network’s indirect connectivity. Constructing the Vietoris-Rips complex (Vietoris 1927) on the PPI network allows counting “non-trivial loops”: loops made of chains of PPIs (the PPIs are the edges of our Vietoris-Rips complex), such that some of the proteins involved were not directly connected to each other by a PPI. A pair of such proteins was then indirectly connected by two chains of PPIs along the loop (each obtained by starting from one protein and following the loop in one of the two directions until reaching the protein which was not directly connected to it by a PPI). For this reason, we called a non-trivial loop an “indirect connection”.

The number of indirect connections and the number of PPIs are positively correlated, and previously we observed that the ratio of these two (number of indirect connections divided by number of PPIs) varies only moderately when we varied the StringDB combined score threshold, which modified the PPI network topology by reducing the number of PPIs (Decano et al 2020). If the indirect connectivity of the PPI network was defined as this ratio for one score threshold, it would lose information about the strength of the PPIs. Therefore, we introduced here a measurement which takes the PPI network topologies at all thresholds into account; we called it the Persistent Maximum of Non-trivial Loops per Edge (PMNLE). This is the maximal ratio (number of indirect connections divided by number of PPIs) which was reached or exceeded across an interval of at least 100 score thresholds. Measuring the PMNLE by calculating the indirect connections at all score thresholds was computationally inefficient, so to reduce the ecological impact and computation time 20-fold, we used an approximate PMNLE (aPMNLE), which was the PMNLE measured only at score thresholds from 400 to 900 with a step of 20. For 16 samples, the scaled deviation between the PMNLE and aPMNLE was small at just 0.02% with a standard deviation of 0.25% (Table S2).

The PPI networks of the 491 samples with plasmid-encoded proteins were analysed with all proteins, and then with chromosomal ones only. This used a specialised open-source tool (Rahm 2019) that constructed the two-dimensional part of the Vietoris-Rips complex by registering the trios of proteins (namely, triangles consisting of three proteins connected to one another by three PPIs) and then ran sparse matrix computations with the LinBox library (with LinBox v1.1.6) to obtain the 1^st^ Betti number, which is the number of indirect connections, and the 0^th^ Betti number, which is the number of connected components (subnetworks of the PPI network in which each pair of proteins is joined by a chain of PPIs -hence there is no chain of PPIs joining two distinct connected components). Additionally, we use the numbers of PPIs, trios, connected components and indirect connections computed at the combined score threshold 400 for comparison: where unstated, the combined score threshold used was 400. In many bacteria’s PPI networks, the purely chromosomal part determined the indirect connectivity, i.e., if the aPMNLE for the full PPI network was similar to the aPMNLE of the chromosomal PPI network, then the influence of the plasmids on the aPMNLE was negligible. We examined the difference of the former minus the latter of these two aPMNLE values, which we scaled by the aPMNLE of the full PPI network to measure the percentage change when plasmid-linked proteins were removed. The aPMNLE was associated with the numbers of PPIs per sample (r=0.42, p=8e-182).

## Results

We retrieved PPI data from StringDB to compare the properties of plasmid-encoded versus chromosomal proteins, with a view to the potential compatibility and signals of co-evolution for plasmid-chromosome combinations. This focused on the numbers of chromosome-linked PPIs, plasmid-associated PPIs, taxonomical classification based on plasmid gene profiles, relative rate of PPIs, indirect connections, and the effect on PPI network structure.

### A positive correlation between the numbers of proteins and the numbers of interactions

We examined the association between the numbers of proteins and PPIs across representatives spanning 4,363 valid bacterial samples based on their unique annotated proteins on StringDB. As expected, the numbers of proteins and PPIs was highly correlated (adjusted r=0.914, p=5.2e-15) (Figure 2, Table S3 – see full data at FigShare https://doi.org/10.6084/m9.figshare.19525708). This suggested that larger chromosomes have more proteins and thus PPIs. It also suggested variation in protein and PPI numbers across bacteria did not skew the association between the numbers of proteins and PPIs, enabling evaluation of features of plasmid-encoded proteins in different samples combinations in more detail.

**Figure 2.**
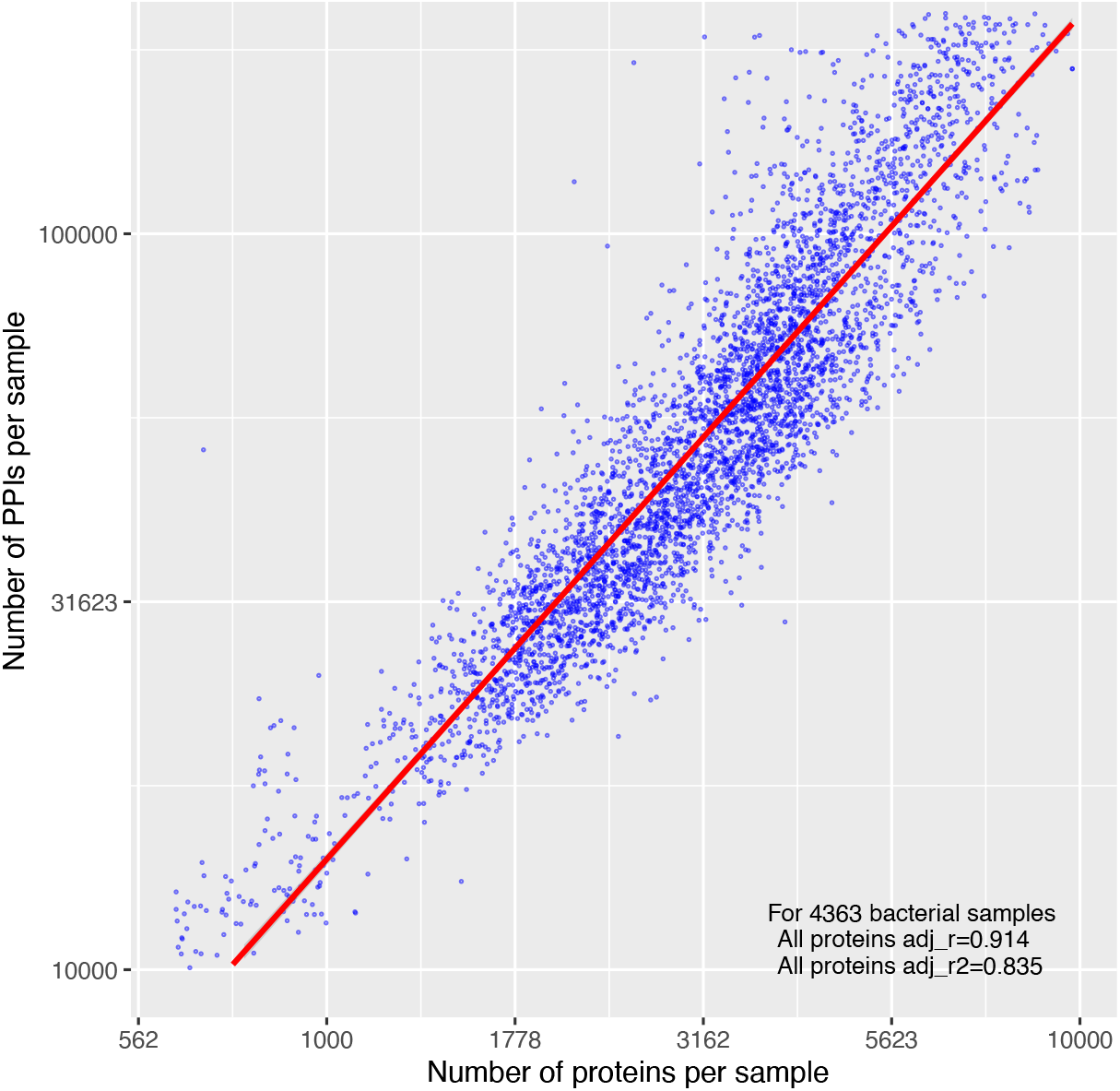
The number of proteins per bacterial genome (x-axis) was highly predictive of the number of PPIs (y-axis) (adjusted r^2^=0.835). The data was for 4,363 bacterial samples, each of which is shown by a blue dot, with a linear model line of best fit shown in red.

### Distinct patterns of plasmid-encoded gene retention across bacterial samples

We compared these 4,363 samples’ PPI data with the set of 8,941 unique plasmid-linked genes where a gene was defined as plasmid-encoded if it was detected on a plasmid from the same species. 66.2% (5,918) of these genes were not detected, yielding 3,023 detected genes. 491 samples (11.2%) had plasmid-encoded genes, leaving 88.8% (3,872) samples with no such genes detected. In this set of 491, the median number of samples in which a plasmid-encoded gene was found was three (interquartile range four, Figure 3A), and the median number of plasmid-encoded genes per sample was five, with an interquartile range of 12 (Figure 3B). The fraction of plasmid-encoded genes per sample was 0.65±2.5% (27±112 out of 3,927±1,367, mean±SD) in these 491 (Figure 3). Six *E. coli* had a median of 1,035 plasmid-encoded genes per sample and were the only samples with over 400 plasmid-encoded genes. These results illustrated that most plasmid-encoded genes were rare, with few common ones, and that widely shared plasmid-encoded genes were unusual (Figure S2, see full figure FigShare doi: https://doi.org/10.6084/m9.figshare.19525453).

**Figure 3.**
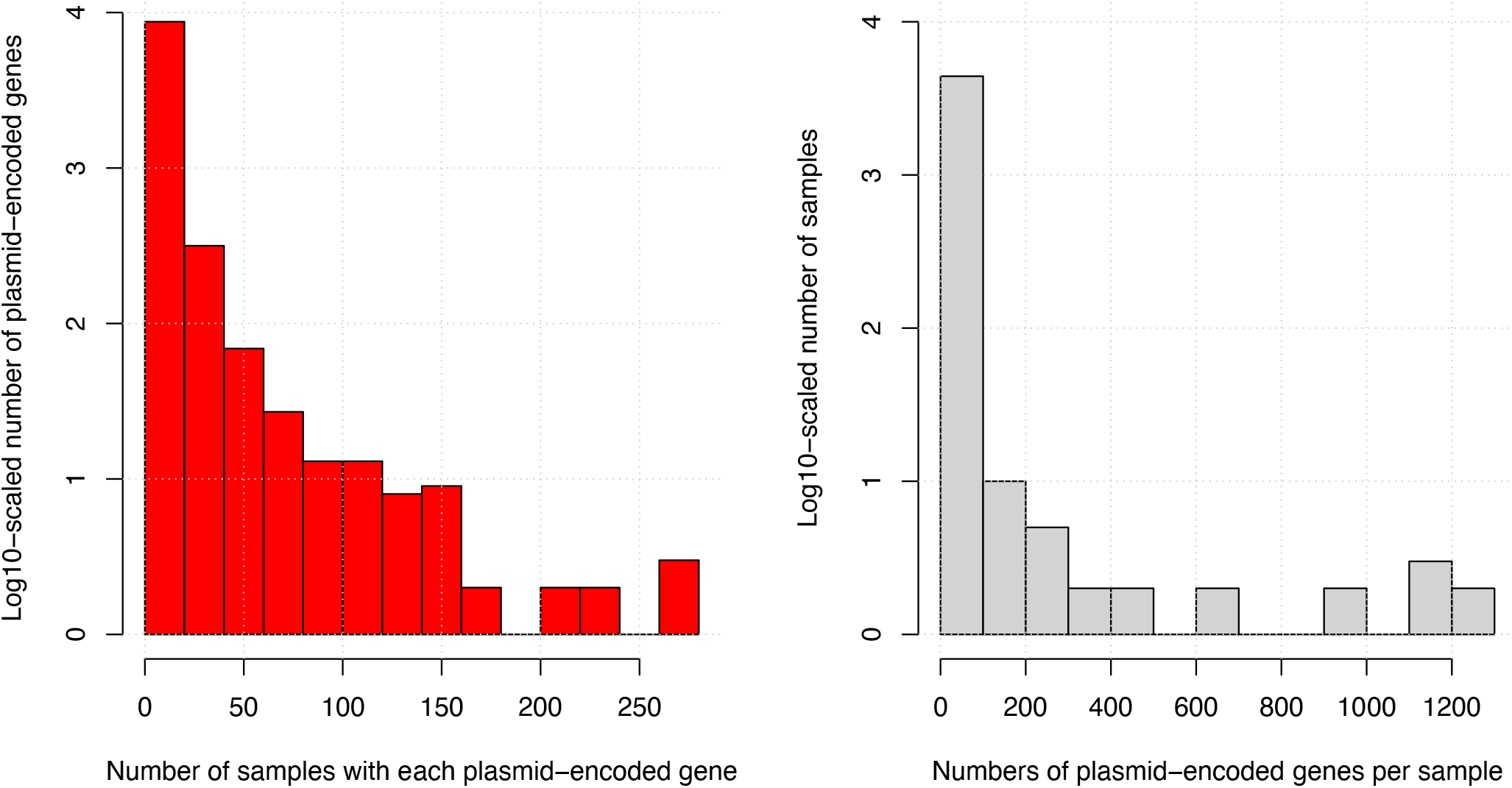
Left: The number of samples in which each plasmid-encoded gene (red) was found was generally small with a median of five. Right: The number of plasmid-encoded genes per sample (grey) had a median of three, including six *E. coli* samples with a median of 1,035 plasmid-encoded genes per sample that represent all samples with >400 genes. These figures excluded samples with no plasmid-encoded genes.

### Plasmid-encoded genes are structured and reflect taxonomic groups

The samples were clustered by similarity based on their binary 3,023 unique plasmid-encoded gene data (presence-absence). Taxonomic classifications across the 100 families, 46 orders, 20 classes and ten phyla in these 491 samples showed that the *Enterobacteriaceae* were the most diverse family (n=24 samples, Z=1.49), and that the *Enterobacterales* (n=57, Z=0.77) was the most variable order, with limited changes at the class and order levels. Overall, the plasmid-encoded gene profiles per taxonomic group did not yield an informative taxonomic resolution (Figure S3, see full figure FigShare doi: https://doi.org/10.6084/m9.figshare.19525435). Evidence for this low overall covariation came from PCA of this data where six *E. coli* and three *Ralstonia solanacearum* samples explained most variation across PC1 (34.5%) and PC2 (5.6%) (Figure S4A). These six *E. coli* had far more plasmid-encoded genes compared to the sample with the next highest number (*Ralstonia solanacearum* PSI07 with 347), which in turn exceeded the other samples substantially. Excluding these nine divergent *E. coli* and *R. solanacearum* samples, the remaining 482 samples’ PC1 had 7.0% of variation and PC2 had 5.0% (Figure S4C). The plasmid-encoded proteins had a similar rate of covariation across all 491 samples where 16.8% of variation was allocated to PC1, and 14.9% to PC2 (Figure S4B), with higher heterogeneity in the 482 samples (6.8% to PC1, 4.9% to PC2, Figure S4D, see full figure at FigShare doi: https://doi.org/10.6084/m9.figshare.19525465, and data at FigShare doi: https://doi.org/10.6084/m9.figshare.19525471). These patterns were largely recapitulated by LSA of all samples, which showed limited variation across the 1^st^ (10.0% of variation) and 2^nd^ (4.1% of variation) dimensions (Figure S5).

### Plasmid-encoded proteins are rare and collectively interact with diverse chromosomal proteins

We examined the samples’ 3,023 unique plasmid-encoded proteins to identify PPIs where one or both proteins were plasmid-related (termed plasmid-related PPIs here, see data at FigShare doi: https://doi.org/10.6084/m9.figshare.19672527) to distinguish them from chromosomal PPIs where no plasmid-encoded proteins were found, and we also tracked the rarer category of plasmid-related PPIs where both protein pairs were plasmid-encoded (see data per sample at shorturl.at/mtFNZ). Overall, as a percentage of all PPIs, plasmid-related PPIs constituted a median of 0.35±4.76% (mean±SD) PPIs per sample, including 0.02% of all PPIs exclusively involving plasmid-encoded proteins. 3,874 samples had zero detected plasmid-related PPIs, 489 samples had at least one plasmid-related PPI (two samples had one plasmid-encoded protein with no PPIs). These 489 samples had a median of 169 PPIs where at least one protein was plasmid-related and a median of 53,947±27,590 chromosomal PPIs per sample, which was comparable to the rate for all samples (48,834±46,615, Table S3). Their total number of plasmid-related PPIs had no clear association with the numbers of proteins per sample (r=0.05), but did with the numbers of chromosomal PPIs per sample (r=0.27, Figure S6). Of these 489 samples, 283 had at least one PPI involving only plasmid-encoded proteins, with a median of five such PPIs.

We examined the plasmid-gene associations across 331 samples that had at least three of the 1,733 plasmid-encoded proteins detected in at least three samples using NMF (Figure S7). This allocated these samples and proteins to six distinct groups based on their pattern of plasmid-encoded protein sharing (Figure S8, see full figure at FigShare doi: https://doi.org/10.6084/m9.figshare.19525492), and the proteins to these same six groups as mixture coefficients based on their prevalence in the samples (Figure S9, see full figure at FigShare doi: https://doi.org/10.6084/m9.figshare.1952559). There were just five samples in rank 3 that were five *E. coli* (all bar *E. coli* 536), which was far below the mean number of samples per rank (77.3, Figure S10). These were associated with 1,065 plasmid-encoded genes, far above the average number of proteins per rank (526.2, Figure S11).

Similarly, we investigated 2,363 plasmid-encoded proteins’ associations based on PPI rates across the same 481 samples where each protein had >30 PPIs and each sample had >10 PPIs. This allocated these samples and proteins to seven distinct groups based on their pattern of PPI numbers (Figure S12, see full figure at FigShare doi: https://doi.org/10.6084/m9.figshare.19611276), and these same proteins to the seven groups as mixture coefficients based on their PPI numbers for each sample (Figure S13, see full figure at FigShare doi: https://doi.org/10.6084/m9.figshare.19611282). Like above, all six *E. coli* (including *E. coli* 536) were the sole members of a rank (five), and corresponded to 238 proteins’ PPI rates as mixture coefficients (Figure S14), which again was more than the average number of PPI rates associated with each rank (748, Figure S15).

### Plasmid-encoded proteins have two-fold higher rates of protein-protein interactions

The mean number of PPIs per protein was 2.2-fold higher for plasmid-related proteins compared to chromosomal ones using data for all samples, though with large variations (n=13,348 plasmid-related proteins in 491 samples: 39.0±35.0 vs n=15,450,151 chromosomal proteins in 4,363 samples: 17.6±5.7, mean±SD, t-test p=4.8e-16, Figure 4). This was not due to artefacts in the 489 samples with plasmid-related PPIs: their rate of chromosomal PPIs at 14.8±3.7 PPIs per protein was similar to the other samples (Table S3, see data at FigShare doi: https://doi.org/10.6084/m9.figshare.19575820).

**Figure 4.**
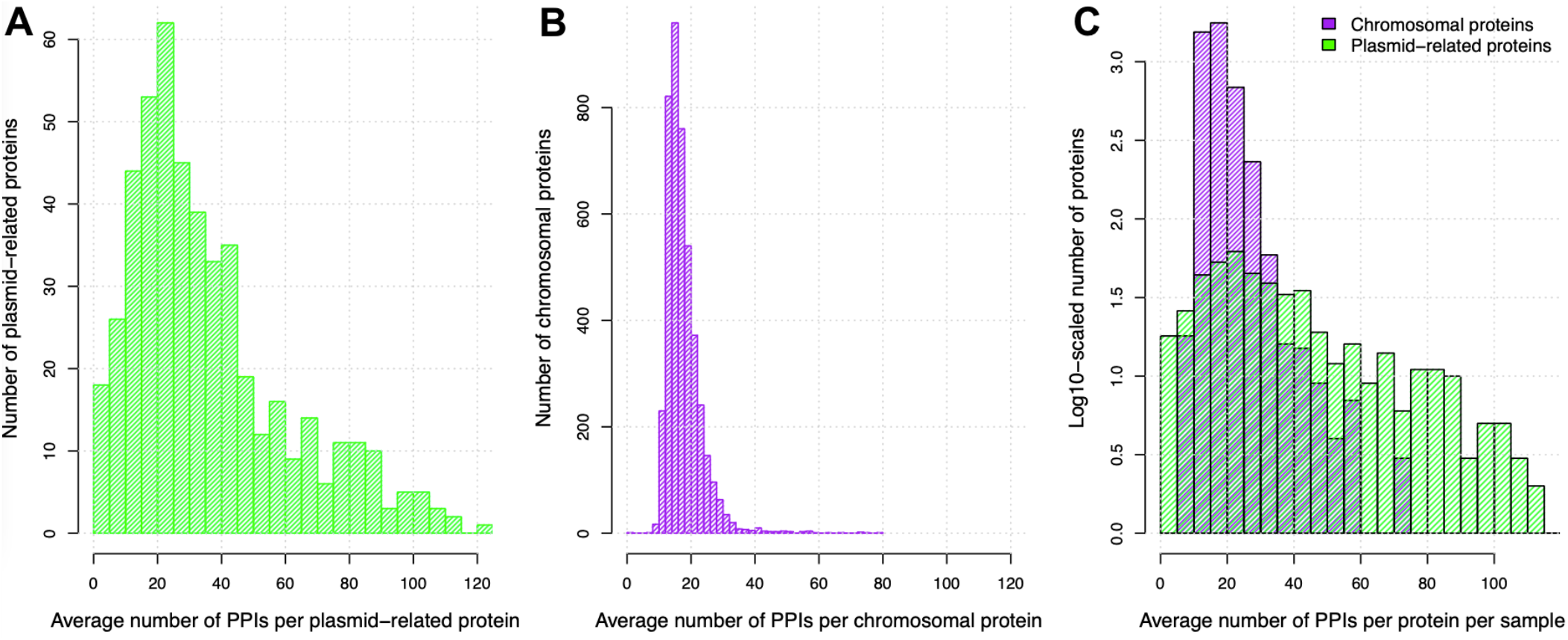
The average number of PPIs per (A) plasmid-related protein (green, 39.0±35.0, mean±SD) was higher than that for (B) chromosomal proteins (purple, 17.6±5.7), but (C) there was no consistent evidence for a consistent difference in PPI rates between these two groups (plasmid-related proteins in green, chromosomal proteins in purple – note log10-scaled y-axis and minor differences in binning per plot).

However, this effect was protein-specific and not consistently discriminatory because these different groups overlapped considerably (Figure 4C), implying that genomic context was not a major determinant of PPI rates, and plasmid-related proteins simply had a wider range of PPI levels. To control for the high variation in PPI rates per plasmid-related protein, we found that 19% (65 out of 345) of samples with sufficient numbers of plasmid-related proteins and PPIs to test quantitatively had plasmid-encoded proteins with much higher PPI rates per sample (t-test p<0.05), including all six *E. coli* (Figure S16, see data at FigShare doi: https://doi.org/10.6084/m9.figshare.20230299). As an illustration of PPI network structure, a network centred on the SfmC protein in *E. coli* K12 MG1655 showed high levels of connectivity amongst 20 other proteins (Figure S17, Table S4).

Nearly all (>99%) plasmid-encoded proteins had a PPI with a chromosomal protein, whereas only 7% of plasmid-encoded proteins had a PPI with another plasmid protein in the 489 samples with at least one plasmid-related PPI. Using the plasmid protein frequency information above, we tested for structure between plasmid-related proteins and chromosomal ones based on the rates of intergroup PPIs in a manner analogous to Wright’s F-statistic to quantify the difference between the observed and expected levels of plasmid-chromosome and plasmid-specific PPIs per sample (Wright 1951). We found a relatively higher rate of plasmid-plasmid PPIs compared to plasmid-chromosome PPIs, which in turn were much higher than chromosomal PPIs (Figure S18). This suggested that plasmid-encoded proteins tended to interact with other plasmid proteins more often than would be expected by chance, indicating PPI network separation of plasmid-encoded and chromosomal proteins. Moreover,

### Protein-protein interaction networks are generally robust to the loss of plasmid proteins

To examine the effect of plasmid-encoded proteins on PPI network topology, we examined the aPMNLE for all proteins and for chromosomal proteins only, such that the difference between these aPMNLE values indicated plasmid-driven effects (Figure S19). The aPMNLE reflects indirect (or secondary) connections, which may be reduced when plasmid-related proteins are removed if there are no alternate paths between remaining chromosomal proteins. However, if there are many PPI paths between chromosomal proteins, then this high indirect connectivity may indicate robustness within the PPI network such that the elimination of plasmid-related proteins has no effect (plasmid-related proteins thus have few indirect connections). Moreover, if plasmid-related proteins tend toward the PPI network periphery, the total number of indirect connections may not change relative to a larger drop in the numbers of PPIs, again indicating plasmid-related proteins had few indirect connections. In contrast, if plasmid-related proteins are central and mixed among chromosomal proteins in the PPI network, the number of indirect connections and numbers of PPIs will be closely correlated (here, plasmid-related proteins have proportional indirect connection rates).

A majority of samples (96.3%, 471 out of 489 samples) showed no substantive differences in aPMNLE values between all proteins versus chromosomal ones alone, indicating that plasmid-encoded proteins had no large effects on the PPI network structure even though the plasmid-related proteins contributed large numbers of PPIs, consistent with robust networks. This was consistent with the idea that for these 471 samples the plasmid-encoded proteins overall had comparable indirect connection rates with the chromosomal proteins, but perhaps higher relative PPI rates because no evidence of excessive indirect connection numbers was observed.

3.7% (18 out of 489) samples had an aPMNLE for all proteins (including plasmid-encoded ones) exceeding the chromosomal aPMNLE by two standard deviations, indicating that plasmid-encoded proteins increased the indirect connection rates and that these PPI networks were less robust. No samples had a chromosomal aPMNLE that was two standard deviations above the full network aPMNLE, so no plasmid-encoded protein set strongly increased the rate of indirect connections per sample. When compared to the other 471 samples with plasmid genes, these 18 had altered aPMNLE values for chromosomal proteins (t-test p=4.9e-5) but not for all proteins (p=0.82, Figure S20). Furthermore, if the plasmid-encoded proteins were included for each sample, the number of PPI network connected components grew by 71% for the 471 samples with robust networks, whereas for these 18 samples with less robust networks it grew by 327%, indicating that these plasmid-encoded proteins had high PPI rates, fewer indirect connections and were often disconnected from the main network. We also observed these 18 had fewer genes per sample compared to the larger set of 471 (2,314±1,429 vs 4,013±1,336, mean±SD), perhaps because smaller genomes have fewer PPIs overall and so were less robust to plasmid-related protein removal.

To quantify the extent to which plasmid-encoded proteins increased PPI rates but not indirect connections, we compared the correlation between plasmid-related proteins and PPIs per sample in the 489 samples with plasmid-related PPIs, which were positively correlated (r=0.51, p=1e-143, Figure 5A). However, increasing the number of plasmid-encoded proteins did not have the same magnitude of effect on the number of indirect connections per sample (r=0.06, p=1, Figure 5B). Similarly, the latter was negatively associated with the number of plasmid-related PPIs (r=-0.08, p=0.30, Figure 5C). The partial correlations for each pair of variables were similar (0.86, 0.04, -0.10, respectively). For example, the 69 plasmid-encoded proteins of *Serratia marcescens* subsp. marcescens Db11 had 92 plasmid-plasmid PPIs with one another and an additional 2,214 plasmid-chromosome PPIs with (an average of 33.4 PPIs per plasmid protein), compared to 58,857 chromosomal PPIs among the 4,614 chromosomal proteins (12.8 PPIs per chromosomal protein, Table S4, Figure S21). Overall, this implied that plasmid-related proteins tended to be at the PPI network periphery connected to chromosomal proteins, and not mixed usually with the main chromosomal proteins in the PPI networks.

**Figure 5.**
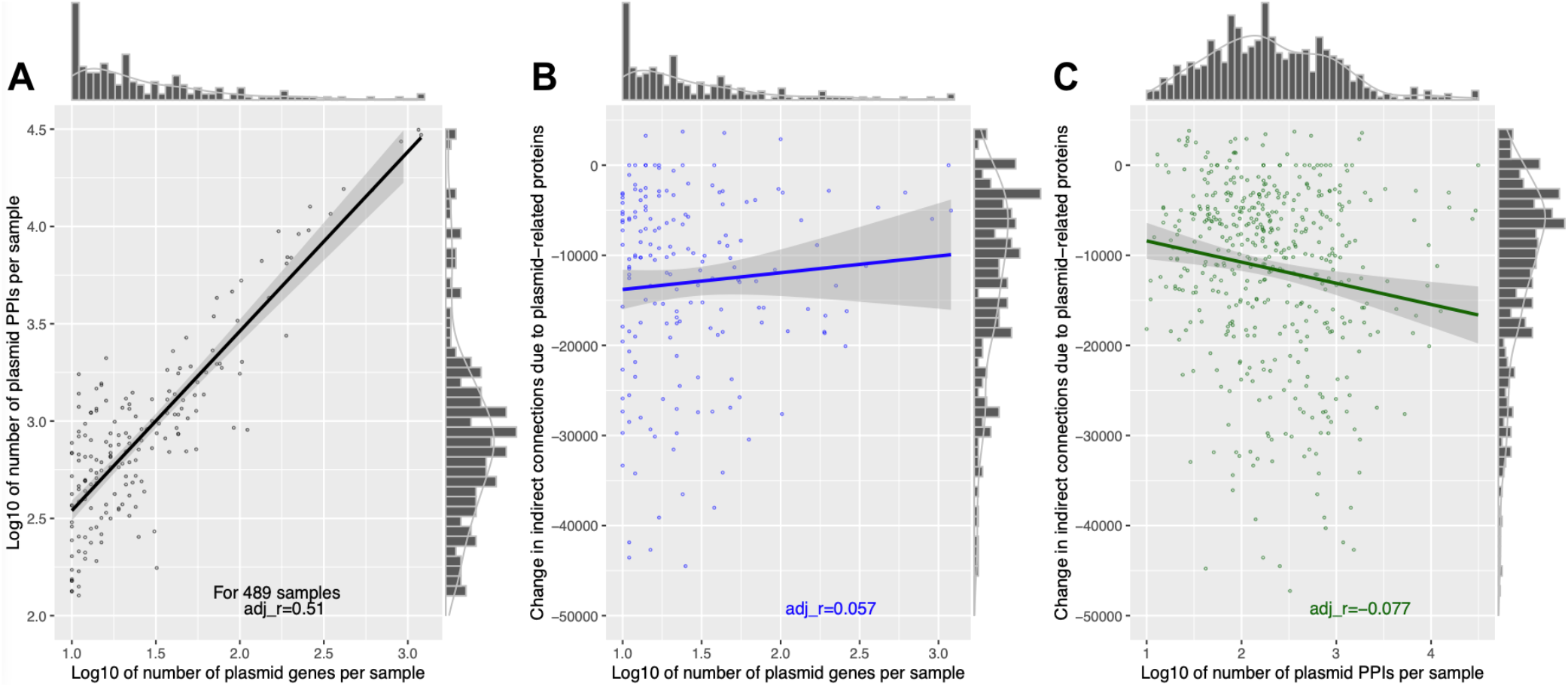
The (A) log10-scaled numbers of plasmid-related proteins per sample (x-axis) was positively correlated with the numbers of plasmid PPIs per sample (y-axis) (black, r=0.51) in the 489 samples with plasmid-related PPIs, but less so with (B) the change in the numbers of indirect connections per sample due to the inclusion of the plasmid-encoded proteins (blue, r=0.06). (C) The log10-scaled numbers of plasmid PPIs per sample (x-axis) was negatively associated with the numbers of indirect connections per sample due to the inclusion of the plasmid-encoded proteins (green, r=-0.08). The lines of best fit for each plot shows the linear correlation of the number of PPIs per sample with the number of indirect connections per protein. The histograms indicate the marginal densities per axis for each dataset.

### Plasmid-related proteins are generally peripheral to PPI networks

Given that plasmid-related proteins purported to be highly connected but not central to PPI networks mainly composed of chromosomal proteins, we scaled the numbers of indirect connections by protein. We observed a trimodal pattern of indirect connections per protein. In the 489 samples with plasmid-related PPIs, the chromosomal proteins had more indirect connections per protein than plasmid-encoded proteins (average per sample 3.7±2.1 vs 0.9±0.6, t-test p=5.2e-15). Moreover, the indirect connections per protein in these 489 was positively correlated with the number of PPIs per sample for chromosomal (r=0.61, Figure 6A) but not all proteins (r=0.02) due to the effect of plasmid-encoded proteins, confirming that the latter tended not to be mixed among the PPI networks. The chromosomal proteins of those 489 samples in turn had a lower level of indirect connections per protein compared to the 3,874 samples without plasmid-related PPIs (average per sample 3.7±2.1 vs 5.5±3.1, t-test p=5.2e-15). The latter 3,874 samples had higher levels of indirect connections that were positively correlated with the PPIs per sample like the 489’s chromosomal proteins (Figure 6B).

**Figure 6.**
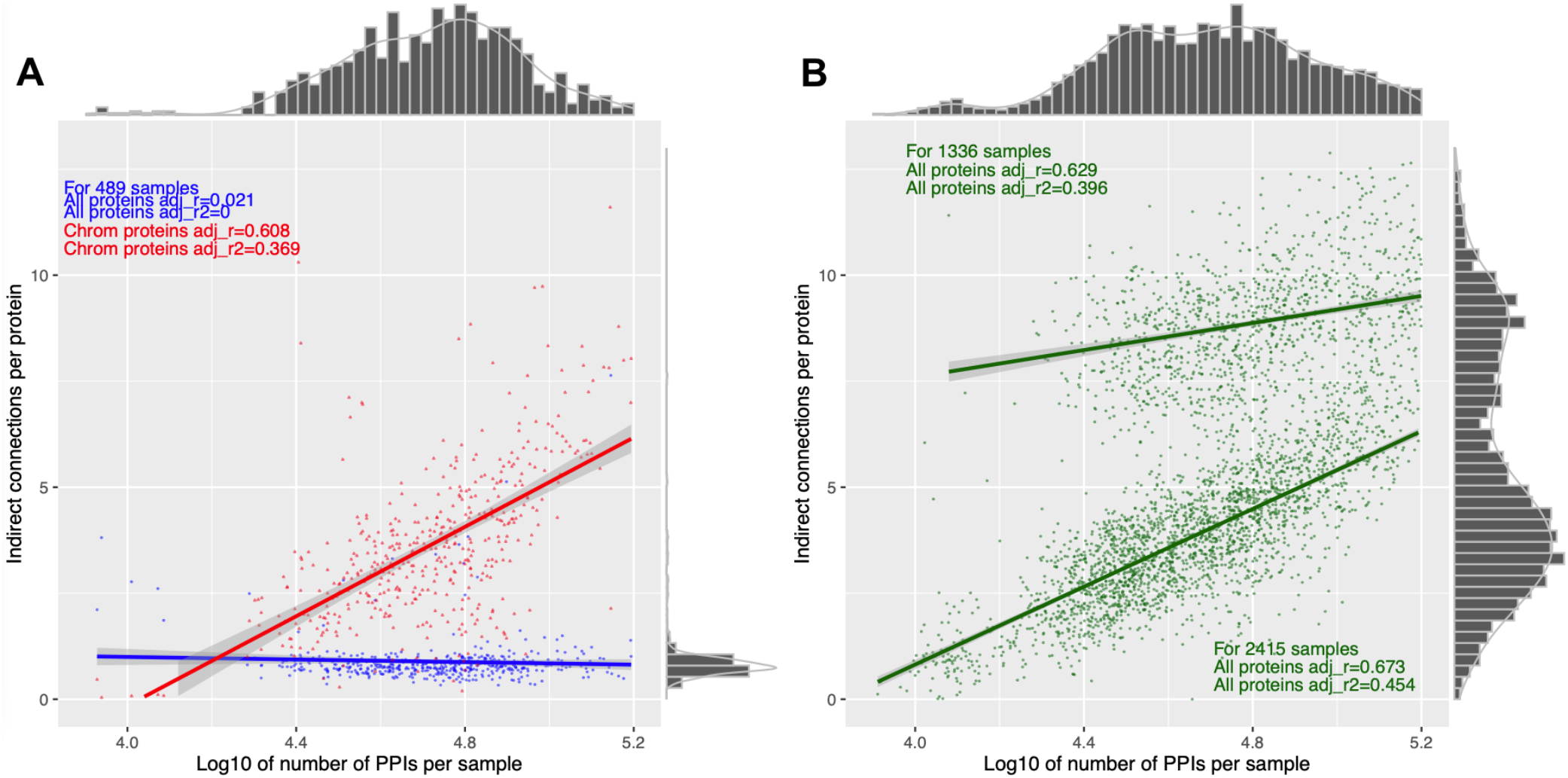
The number of PPIs per sample (x-axis) was positively correlated with the number of indirect connections per protein for (A) chromosomal (red, r=0.61) but not all proteins (including plasmid-encoded ones, blue, r=0.02) in the 489 samples with plasmid-encoded proteins. (B) The other samples with no plasmid-encoded proteins had positive correlations (green, r=0.63 and r=0.67) though with a bimodal pattern (groups of n=2,415 and n=1,336). The lines of best fit for each plot shows the linear correlation of the number of PPIs per sample with the number of indirect connections per protein. The histograms indicate the marginal densities per axis for each dataset.

Plasmid-related proteins had a median of 4.4-fold fewer indirect connections per sample than chromosomal proteins in the set of 489 samples (Figure 7A). We also observed that additional plasmid-chromosome PPIs constituted 89.7% of all PPIs in these 489, and 93% of plasmid-encoded proteins had no plasmid-related PPIs. To explain this, consider a PPI network where part of it had four chromosomal proteins (A, B, C, D) sharing PPIs A-B, B-C, C-D and D-A, so it has one indirect connection (with a path A-B-C-D, Figure S22). If we added a plasmid protein P to this network such that it had PPIs with all four proteins, the indirect connection across A-B-C-D would be lost but the network would gain four PPIs. An alternative model where A, B, C and D had no PPIs with one another would be less likely because then the addition of P would create PPIs and more indirect connections, which was not observed in our findings. In addition, if A, B, C and D interact with P then it is more likely that they are both functionally related and interact with one another.

**Figure 7.**
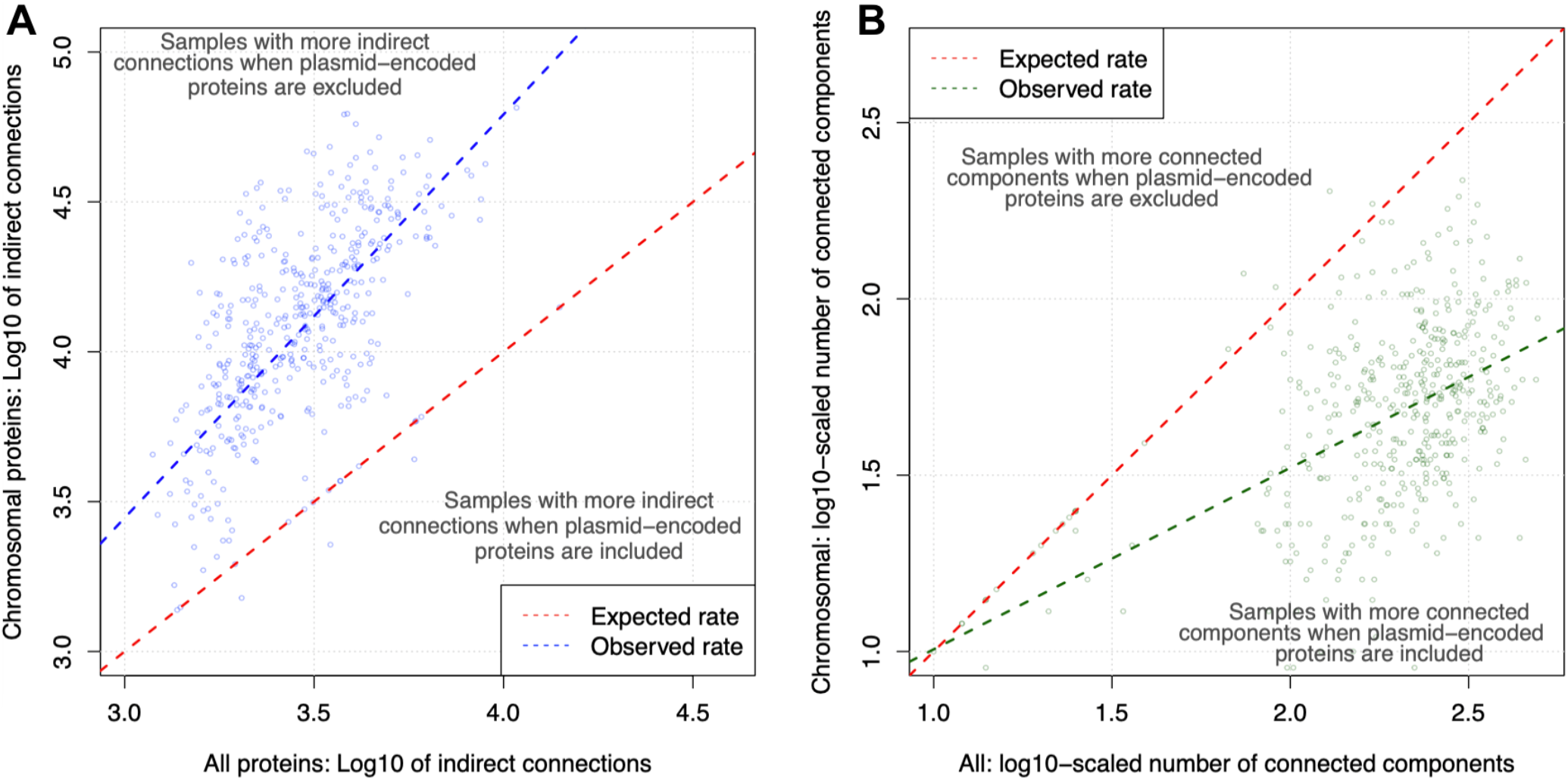
The log10-scaled numbers of (A) indirect connections (blue) and (B) connected components (green) for all (chromosomal and plasmid) proteins (x-axis) related to chromosomal proteins only (y-axis) for the 489 samples with plasmid-related PPIs. The red lines show the expected rate if (A) the numbers of indirect connections for all proteins and chromosomal proteins were identical and (B) the numbers of connected components for all proteins and chromosomal proteins were identical. (A) The blue line shows the correlation between the numbers of indirect connections for all proteins and chromosomal proteins (>98% of samples had more indirect connections when plasmid-related proteins were omitted). (B) The green line shows the correlation between the numbers of connected components for all proteins and chromosomal proteins (>97% of samples had more connected components when plasmid-related proteins were retained).

Finally, we assessed the numbers of PPI network connected components for all proteins versus chromosomal ones: the latter had fewer connected components compared to all proteins in the 489 samples with plasmid-related PPIs (50.0±34.6 vs 227±95.9, median±SD, Figure 7B). The numbers of proteins per connected component was more strongly associated with the chromosomal proteins than for all proteins in relation to the numbers of PPIs (r=0.35 vs 0.01), chromosome-related PPIs (r=0.35 vs 0.00), and proteins per sample (r=0.18 vs 0.00) but not plasmid-related PPIs (r=0.04 vs 0.00, Figure S23) in these 489. Additionally, these 489 samples had far more connected components compared to 3,872 samples without plasmid-related PPIs (means 222.4 vs 40.5, t-test p=4.8e-15). This highlighted that when there were more PPIs and proteins present, chromosomal proteins were spread across fewer connected components, whereas plasmid proteins could be found in more separate connected components.

## Discussion

Bacterial genome structure has been shaped by HGT more so than gene duplication in many species, such as *E. coli* (Lercher & Pál 2008, Price et al 2008). Mobile plasmid-encoded genes can endow host cells with new phenotypes, including those linked to changed adhesion, AMR and virulence (Lipworth et al 2022), and their fitness effects are associated with specific genetic, regulatory and protein interactions (Hall et al 2017). To further our understanding of how new plasmid-encoded proteins can affect the structure and signalling bacterial PPI networks, we compared the PPI profiles of plasmid-linked and chromosomal proteins across 4,363 representative samples of well-characterised bacteria using data from StringDB to find 491 samples with evidence of plasmid-encoded genes within the same species.

We found that the number of proteins per sample was correlated with the number of PPIs, as anticipated. This allowed further exploration to compare plasmid-encoded proteins with chromosomal ones. Most plasmid-encoded proteins were rare across bacteria, and the total fraction of the genome they constituted varied extensively. We defined plasmid-related PPIs as PPIs where at least one of the interacting proteins had previously been observed on a contig labelled as a plasmid within the same species, separate from chromosomal PPIs where neither protein was plasmid-linked. Only 11.2% of samples (489 out of 4,363) had plasmid-related proteins and PPIs. Even in these samples, plasmid-related genes made up only 0.65% of all genes detected and thus were rare. As a consequence, plasmid-related PPIs made up only 0.14% of all PPIs, and PPIs that were exclusively plasmid-related were very rare (0.016%).

Plasmid-encoded proteins covaried with one another depending on the genetic background of the host cell such that the clustering the samples based on their plasmid-encoded gene similarity identified six groups of covarying plasmid-encoded proteins and bacterial samples, including a specific one for *E. coli* alone. This aligned with existing expectations of plasmid-host co-evolution (Loftie-Eaton et al 2017, Jordt et al 2020). The *Enterobacterales* was the clearly the most variable order, but overall known taxonomical differences were not apparent. Similarly, the PPI rates per protein per sample had even less taxonomical resolution power, showing that the numbers of PPIs per protein was typically a more dynamic feature.

The complexity hypothesis (Jain et al 1999) asserts that proteins undergoing HGT have fewer PPIs and operational cellular processes rather than not informational ones like transcription or translation (Rivera et al 1998, Jain et al 1999, Jain et al 2002). It has been asserted that proteins with many PPIs are less likely to undergo HGT (Wellner et al 2007, Price et al 2008, Cohen et al 2011), but we found that plasmid-linked proteins had two-fold more PPIs on average. Nonetheless, the predictive power of this across chromosomal versus plasmid-related proteins was low, suggesting that there is no intrinsic difference in PPI rates between these groups of proteins. Nuances to the complexity hypothesis have been found before: firstly, the being in a protein complex versus HGT chances (Wellner et al 2007); secondly, the effect of cellular component state, function complexity and function conservation on adaptive evolution potential (Aris-Brosou 2005); thirdly, HGT-linked genes have higher numbers of regulatory factors (Price et al 2008); and fourthly, PPI connectivity has a large effect for proteins with a similar function (Cohen et al 2011). Notably, many previous studies have focused on conserved orthologs, whose genomic contexts and features may differ from genes that are more mobile. Consequently, operational versus informational processes, complex protein functions, an intracellular location, an ancient essential function, and host-recipient genetic distance may be more dominant factors in determining PPI rates than HGT given our results here. An important caveat is that the general trends we observed may not hold for all genera, families or orders.

We found that plasmid-related proteins interacted with large numbers of other chromosomal and plasmid proteins. However, plasmid-related proteins contributed much less indirect (aka secondary) connections, unlike chromosomal proteins. We found evidence of PPI network structure between chromosomal and plasmid-encoded proteins, finding that plasmid proteins had a higher rate of plasmid rather than chromosomal protein partners than expected. In addition, by using the aPMNLE as a measure of indirect connections, most samples with plasmid-related PPIs (>96%) were robust to the loss of plasmid-encoded proteins. The samples with networks more affected by plasmid-related protein loss had smaller genomes, and thus less redundancy in PPI network paths. Altogether, this was consistent with existing assertions that plasmid-encoded proteins tend to be at the periphery of a PPI network (Lercher & Pál 2008, Price et al 2008), perhaps because plasmids seldom encode essential proteins (Wein et al 2021), though gene essentiality depends on the host (Rousset et al 2021).

Our findings were limited by a number of issues. Firstly, our analysis was based on gene name similarity with the assumption that the redundancy in the annotated plasmid gene names would help alleviate the imprecision in gene naming systems, but this did not mitigate fully the limitation that some proteins were likely misclassified as chromosomal or plasmid-encoded. Secondly, many plasmids and bacterial samples are not annotated consistently, so many gene name matches were undoubtedly missed. Thirdly, we assumed that previous studies’ labelling of large contigs as plasmids was correct: we made this (and all other) data accessible so that revisions to plasmid annotation could be made in light of the high number of genes (>500) in certain plasmids (see Table S1): sequence-based plasmid detection tools could help with this task. Fourthly, we obtained gene name data per sample from StringDB (Szklarczyk et al 2018, Szklarczyk et al 2021), and yet 5,918 plasmid-encoded genes were not detected in these samples, which highlighted imprecision either in the plasmid annotation, the sample annotation or (most likely) both. Fifthly, we did not perform sequence alignment nor read mapping to confirm at the sequence level matches, which is a next step following this work. Future work is needed to improve plasmid annotation, genome annotation, contig classification as plasmid or chromosomal and conduct gene comparisons at the sequence level. Future investigations of the effects of plasmid-encoded and chromosomal proteins on PPI networks should focus on specific samples that have known plasmid compositions.

## Acknowledgements

The authors thanks Min Jie Lee for certain R code contributions.

## Author Contributions

TD – Conceptualization, Methodology, Software, Formal analysis, Investigation, Data Curation, Writing - Original Draft, Writing - Review & Editing, Visualisation, Project administration. AR – Methodology, Software, Formal analysis, Writing - Review & Editing. All authors reviewed the manuscript.

## Data Availability Statement

**Data Availability Statement:**
| Item | Brief description | FigShare link |
| --- | --- | --- |
| Table S1 | Plasmid-encoded genes and their associated plasmid link(s) | https://doi.org/10.6084/m9.figshare.19525630 |
| Table S3 | Key features of the bacterial samples | https://doi.org/10.6084/m9.figshare.19525708 |
| Figure S2 | Heatmap of the bacterial plasmid-encoded genes | https://doi.org/10.6084/m9.figshare.19525453 |
| Figure S3 | Dendrogram of the bacteria based on plasmid-encoded gene similarity | https://doi.org/10.6084/m9.figshare.19525435 |
| Figure S5 | Heatmap of the bacteria and plasmid-encoded genes alphabetically sorted | https://doi.org/10.6084/m9.figshare.19525465 |
| Data for Figure S5 | Presence-absence data with bacteria as rows and genes as columns | https://doi.org/10.6084/m9.figshare.19525471 |
| Figure S8 | NMF heatmap of the metasample clustering of plasmid genes | https://doi.org/10.6084/m9.figshare.19525492 |
| Figure S9 | NMF heatmap of the mixture coefficient for the plasmid-encoded genes | https://doi.org/10.6084/m9.figshare.19525591 |
| Figure S12 | NMF heatmap of the metasample clustering of plasmid PPI rates | https://doi.org/10.6084/m9.figshare.19611276 |
| Figure S13 | NMF heatmap of the mixture coefficient for the plasmid-related PPI rates | https://doi.org/10.6084/m9.figshare.19611282 |
| Data for all PPI analysis | 4,363 samples' CSV files for all proteins with gene name, plasmid status and the total number of PPIs per sample | https://doi.org/10.6084/m9.figshare.19672527 |
| Data for chrom PPI analysis | 4,363 samples' CSV files for each sample showing PPIs, associated genes and StringDB identifiers | shorturl.at/mtFNZ |
| Data for Figure S16 | The numbers of chromosomal & plasmid genes for 343 as PDF image files. | https://doi.org/10.6084/m9.figshare.20230299 |

Large datasets available from this study on FigShare, showing the item, description and the corresponding DOI link. See the Supplementary Data for more. The code associated with this paper is available on Github at https://github.com/downingtim/mobilome_2022 and https://github.com/arahm/HomologyLive and data files are there at https://figshare.com/projects/Bacteria_mobilome_2022/136720.

