## Supplementary material for "Bacterial plasmid-associated and chromosomal proteins have fundamentally different properties in protein interaction networks": SupplData


| Page | Item | Description |
|---|---|---|
| 2 | Table S1 | Each of the plasmid-encoded genes and their associated plasmid link(s) across the plasmid accessions. |
| 2 | Table S2 | The complete PMNLE and approximate PMNLE (aPMNLE) values for 491 samples' PPI data from StringDB across all proteins. |
| 2 | Table S3 | Key features of the bacterial samples. |
| 3 | Table S4 | The 21 genes encoding proteins in the PPI network in Figure S18 along with their genomic context (plasmid or chromosome). |
| 4 | Table S5 | Samples (18, 4% of total) with a much higher aPMNLE for all proteins than the chromosomal ones alone. |
| 5 | Figure S1 | The number of long gene names compared to the number of short gene names per sample. |
| 6 | Figure S2 | Heatmap visualising plasmid-encoded genes for the bacterial samples (y-axis) across 3,023 unique plasmid-encoded genes (x-axis). |
| 7 | Figure S3 | Dendrogram showing the similarity across families based on the samples' plasmid gene profiles. |
| 8 | Figure S4 | PCA of the whole dataset showing the variation of: (A) the samples (excluding *E. coli*) across the plasmid-encoded genes found in these samples, and (B) these genes across these samples. |
| 9 | Figure S5 | Visualisation of the $1^{st}$ (x-axis, 10.0% of variation) and $2^{nd}$ (y-axis) dimensions from LSA. |
| 10 | Figure S6 | A positive correlation between the numbers of plasmid- and chromosome-related (x-axis) PPIs. |
| 11 | Figure S7 | Heatmap visualising plasmid interactomes of 491 bacterial samples (y-axis) across 3,023 unique plasmid-encoded genes using NMF (x-axis). |
| 12 | Figure S8 | Heatmap of the basis coefficient matrices, reflecting the chance of allocating 331 samples (y-axis) to six different groups (x-axis) based on their plasmid gene profiles using NMF. |
| 13 | Figure S9 | Heatmap of the mixture coefficient matrices, reflecting the probability of allocation of 1,733 plasmid-encoded genes (x-axis) to six different groups (y-axis) based on their prevalence in 331 samples using NMF. |
| 14 | Figure S10 | The frequency distributions of the rate of allocation of the 331 samples to each of the six ranks from NMF. |
| 15 | Figure S11 | The frequency distributions of the rate of allocation of the 1,733 plasmid-encoded genes to each of the six ranks from NMF. |
| 16 | Figure S12 | Heatmap of the basis coefficient matrices, reflecting the probability of allocation of the 481 samples (y-axis) to seven different groups (x-axis) based on their plasmid PPI rates using NMF. |
| 17 | Figure S13 | Heatmap of the mixture coefficient matrices, reflecting the probability of allocation of 2,363 plasmid-encoded genes to seven different groups based on their PPI rates in 481 samples. |
| 18 | Figure S14 | The frequency distributions of the rate of allocation of the 481 samples to each of the seven ranks from NMF of the plasmid PPI rates. |
| 19 | Figure S15 | The frequency distributions of the rate of allocation of the 2,363 plasmid-encoded genes to each of the six ranks from NMF of the plasmid PPI rates. |
| 20 | Figure S16 | The numbers of chromosomal and plasmid interactions per gene for the six *E. coli* samples. |
| 21 | Figure S17 | An *E. coli* K12 MG1655 (String ID 511145) PPI network from the StringDB website of 21 proteins in Table S4 centred on SfmC. |
| 22 | Figure S18 | The F values for plasmid-plasmid versus plasmid-chromosome PPIs. |
| 23 | Figure S19 | The distribution of aPMNLE values for all proteins, chromosomal ones and the scaled difference between all and chromosomal proteins across the bacterial samples. |
| 24 | Figure S20 | The association between the aPMNLE values for all proteins (x-axis) compared to those for chromosomal proteins (y-axis). |
| 25 | Figure S21 | The plasmid-encoded proteins' PPI network for *Serratia marcescens* subsp. marcescens Db11. |
| 26 | Figure S22 | A model showing how PPI rates can increase but indirect connection numbers can fall. |
| 27 | Figure S23 | Varying levels of a positive correlation with the number of proteins per connected component. |

**Table S1**. Each of the plasmid-encoded genes and their corresponding sample, resulting in 363,051 gene-sample associations in total. See full table FigShare doi: https://doi.org/10.6084/m9.figshare.19525453
allocated to rank 5.

| Sample | PMNLE | aPMNLE | %difference |
|---|---|---|---|
| *Bifidobacterium thermophilum* | 0.5105 | 0.5112 | -0.15% |
| *Bifidobacterium cuniculi* | 0.4935 | 0.4934 | 0.00% |
| *Acinetobacter baumannii* | 0.5213 | 0.5218 | -0.10% |
| *Bifidobacterium choerinum* | 0.5295 | 0.5296 | -0.02% |
| *Bifidobacterium coryneforme* | 0.5281 | 0.5282 | -0.02% |
| *Bifidobacterium magnum* | 0.5379 | 0.5420 | -0.77% |
| *Bifidobacterium minimum* | 0.4948 | 0.4948 | -0.01% |
| *Enterobacter aerogenes* | 0.5190 | 0.5186 | 0.07% |
| *Klebsiella oxytoca* | 0.5422 | 0.5417 | 0.10% |
| *Klebsiella pneumoniae* | 0.5435 | 0.5412 | 0.42% |
| *E. coli* 536 | 0.5639 | 0.5626 | 0.22% |
| *E. coli* ATCC8739 | 0.5461 | 0.5455 | 0.10% |
| *E. coli* BL21DE3 | 0.5706 | 0.5700 | 0.11% |
| *E. coli* CFT073 | 0.5554 | 0.5545 | 0.15% |
| *E. coli* K12 | 0.5907 | 0.5901 | 0.10% |
| *E. coli* O157H7 | 0.5659 | 0.5654 | 0.09% |
| Mean | 0.5383 | 0.5382 | 0.02% |
| Standard deviation | 0.0272 | 0.0269 | 0.25% |

**Table S2**. The complete PMNLE and approximate PMNLE (aPMNLE) values for 16 samples' PPI data from StringDB across all proteins. The scaled mean difference (%difference) between the complete and approximate PMNLE values was small (0.02% with a standard deviation of 0.25%).

**Table S3**. Key features of the bacterial samples. Fraction is the fraction of all genes that were plasmid-encoded. Plasmid_Plasmid_PPI is the number of PPIs exclusively between plasmid-encoded proteins. Plasmid_PPI is the number of PPIs between a chromosomal protein and a plasmid protein. Chrom_PPI is the number of PPIs between chromosomal proteins. PPIs is the total number of PPIs. CTri indicates the number of chromosomal protein trios. Loops indicates the number of indirect connections for chromosomal proteins. Ccs indicates the number of connected components for chromosomal proteins. CTri_all indicates the number of all protein trios. Loops_all indicates the number of indirect connections for all proteins. Ccs_all indicates the number of connected components for all proteins. APMNLE indicates the chromosomal proteins' aPMNLE (indirect connectivity). APMNLE_all indicates the aPMNLE for all proteins. DiffPMNLE stands for the % difference in aPMNLE when plasmid proteins are added to the chromosomal ones. Proteins_ccs_all is the number of connected components per protein for all proteins. Proteins_ccs_chrom is the number of connected components per protein for chromosomal proteins. Indirect_all is the number of indirect connections per protein for all proteins. indirect_chrom is the number of indirect connections per protein for chromosomal proteins. See FigShare doi: https://doi.org/10.6084/m9.figshare.19525708.

| Gene | Context | Number of PPIs |
|---|---|---|
| *atpA* (aka *papA*) | Chromosome | 220 |
| *atpB* (aka *papD*) | Chromosome | 116 |
| *atpC* (aka *papG*) | Chromosome | 105 |
| *atpD* (aka *papB*) | Chromosome | 166 |
| *atpE* (aka *papH*) | Chromosome | 102 |
| *atpF* (aka *papF*) | Chromosome | 132 |
| *atpG* (aka *papC*) | Chromosome | 172 |
| *atpH* (aka *papE*) | Chromosome | 143 |
| *fimA* | Plasmid | 63 |
| *fimC* | Plasmid | 48 |
| *fimD* | Plasmid | 27 |
| *fimF* | Plasmid | 40 |
| *fimG* | Plasmid | 21 |
| *fimH* | Plasmid | 63 |
| *fimI* | Plasmid | 31 |
| *sfmA* | Chromosome | 58 |
| *sfmC* | Chromosome | 44 |
| *sfmD* | Chromosome | 28 |
| *sfmF* | Chromosome | 60 |
| *sfmH* | Chromosome | 71 |
| *yfaL* | Chromosome | 132 |

**Table S4**. The 21 genes encoding proteins in the PPI network in Figure S18 along with their genomic context (plasmid or chromosome) and the number of PPIs per protein across *E. coli* K12 MG1655's entire set of proteins. In this example, the chromosomal proteins had a higher median PPIs per protein than the plasmid-linked ones (110.4 vs 44.0), and notably both chromosomal and plasmid proteins had high variations in the numbers of PPIs per protein (standard deviations 54.8 and 16.9, respectively).

| Sample | Plasmid genes | Chrom genes | aPMNLE all | aPMNLE chr | diff PMNLE | Plasmid PPIs | Chrom PPIs | PPIs |
|---|---|---|---|---|---|---|---|---|
| *Actinobacillus pleuropneumoniae* serovar 5b str L20 | 1 | 1,980 | 0.546 | 0.451 | 0.175 | 25 | 27,494 | 27,519 |
| *Buchnera aphidicola* BCc | 4 | 357 | 0.516 | 0.122 | 0.763 | 87 | 8,598 | 8,685 |
| *Buchnera aphidicola* str Bp | 4 | 501 | 0.546 | 0.231 | 0.578 | 83 | 10,144 | 10,227 |
| *Buchnera aphidicola* str G002 | 4 | 568 | 0.551 | 0.371 | 0.327 | 85 | 11,725 | 11,810 |
| *Candidatus Riesia* pediculicola USDA | 2 | 532 | 0.542 | 0.343 | 0.367 | 42 | 8,431 | 8,473 |
| *Citrobacter rodentium* ICC168 | 1 | 4,745 | 0.545 | 0.451 | 0.172 | 44 | 72,468 | 72,512 |
| *Helicobacter cinaedi* CCUG 18818 | 1 | 2,300 | 0.513 | 0.388 | 0.244 | 38 | 24,825 | 24,863 |
| *Lactobacillus amylolyticus* DSM 11664 | 17 | 1,648 | 0.516 | 0.429 | 0.169 | 580 | 18,667 | 19,247 |
| *Legionella sainthelensi* ATCC 35248 | 3 | 3,383 | 0.535 | 0.447 | 0.163 | 71 | 33,637 | 33,708 |
| *Morganella morganii* subsp morganii KT | 4 | 3,477 | 0.547 | 0.453 | 0.172 | 304 | 48,504 | 48,808 |
| *Pasteurella multocida* | 1 | 2,027 | 0.539 | 0.439 | 0.185 | 45 | 28,209 | 28,254 |
| *Saprospira grandis* str Lewin | 10 | 4,049 | 0.534 | 0.430 | 0.194 | 683 | 29,006 | 29,689 |
| *Serratia marcescens* subsp marcescens Db11 | 69 | 4,614 | 0.556 | 0.456 | 0.178 | 2,306 | 58,857 | 61,163 |
| *Serratia symbiotica* str Cinara cedri | 1 | 666 | 0.616 | 0.207 | 0.664 | 13 | 12,175 | 12,188 |
| *Simkania negevensis* Z | 6 | 2,433 | 0.505 | 0.398 | 0.213 | 77 | 20,673 | 20,750 |
| *Staphylococcus equorum* subsp equorum Mu2 | 9 | 2,699 | 0.562 | 0.441 | 0.216 | 180 | 31,039 | 31,219 |
| *Staphylococcus pseudintermedius* ED99 | 1 | 2,305 | 0.541 | 0.454 | 0.160 | 115 | 28,751 | 28,866 |
| *Yersinia ruckeri* | 3 | 3,234 | 0.531 | 0.443 | 0.166 | 17 | 44,817 | 44,834 |

**Table S5**. The properties of the 18 bacterial samples (18 out 489 samples with >1 plasmid-related PPI, 3.7%) with a much higher aPMNLE for all proteins than the chromosomal ones alone. Chrom stands for chromosomal. "Diff PMNLE" stands for the fraction of difference between the PMNLE for all (chromosomal and plasmid) proteins versus that for chromosomal proteins. In the sample here with the highest rate of plasmid-encoded genes, *Serratia marcescens* subsp marcescens Db11, the plasmid-encoded genes had 33.4 PPIs per protein, whereas chromosomal genes had 12.8 PPIs per protein. The median scaled PPIs per protein in the plasmid-encoded genes per sample had a median 109% higher than that for chromosomal proteins in these 18 samples.

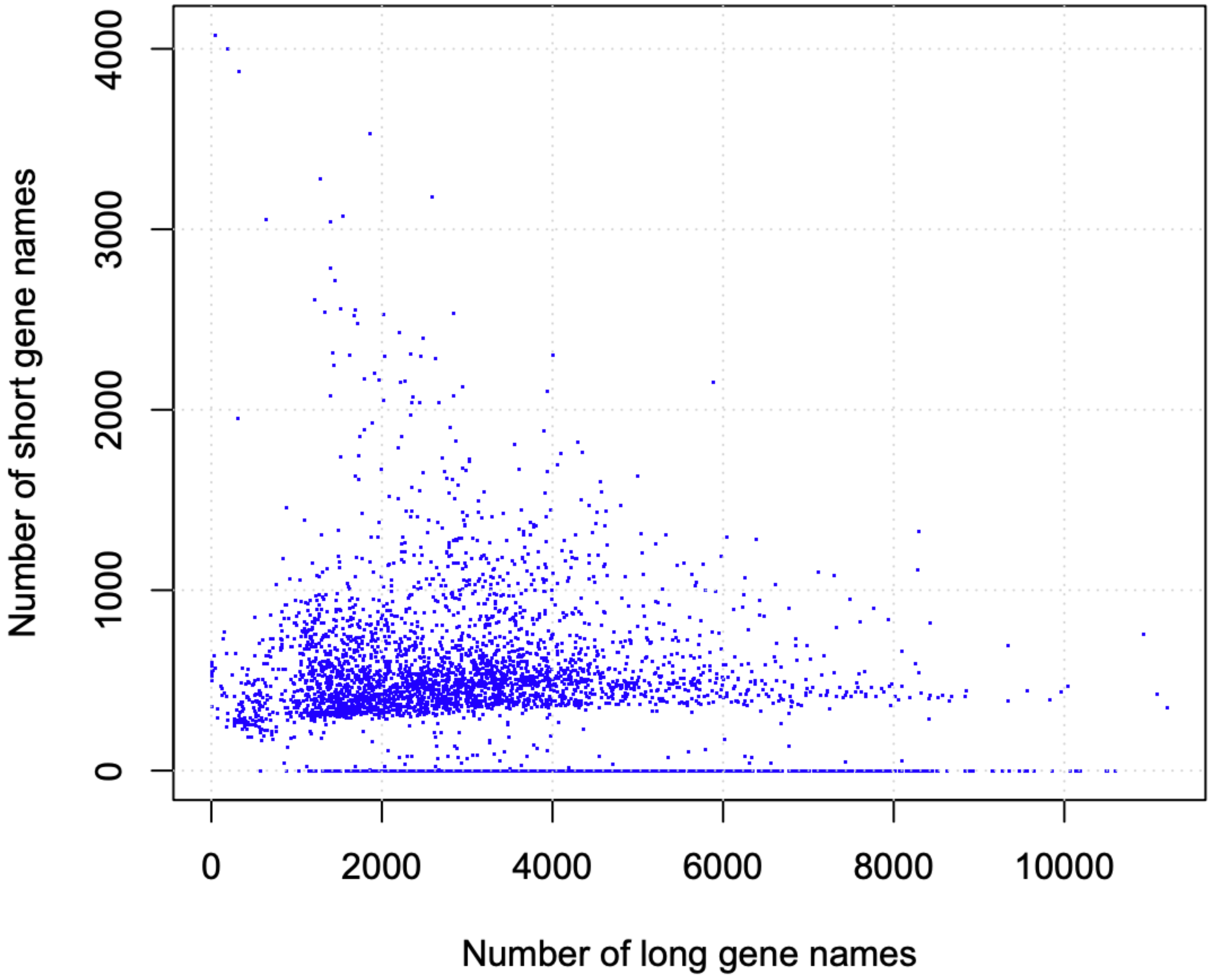


**Figure S1**. The number of long gene names (x-axis) compared to the number of short gene names (y-axis) per sample. The 4,429 samples had a median of 411±410 short names and a median of 2,931±1,744 long names each. Short gene names had four letters or less, and long ones had more than this.

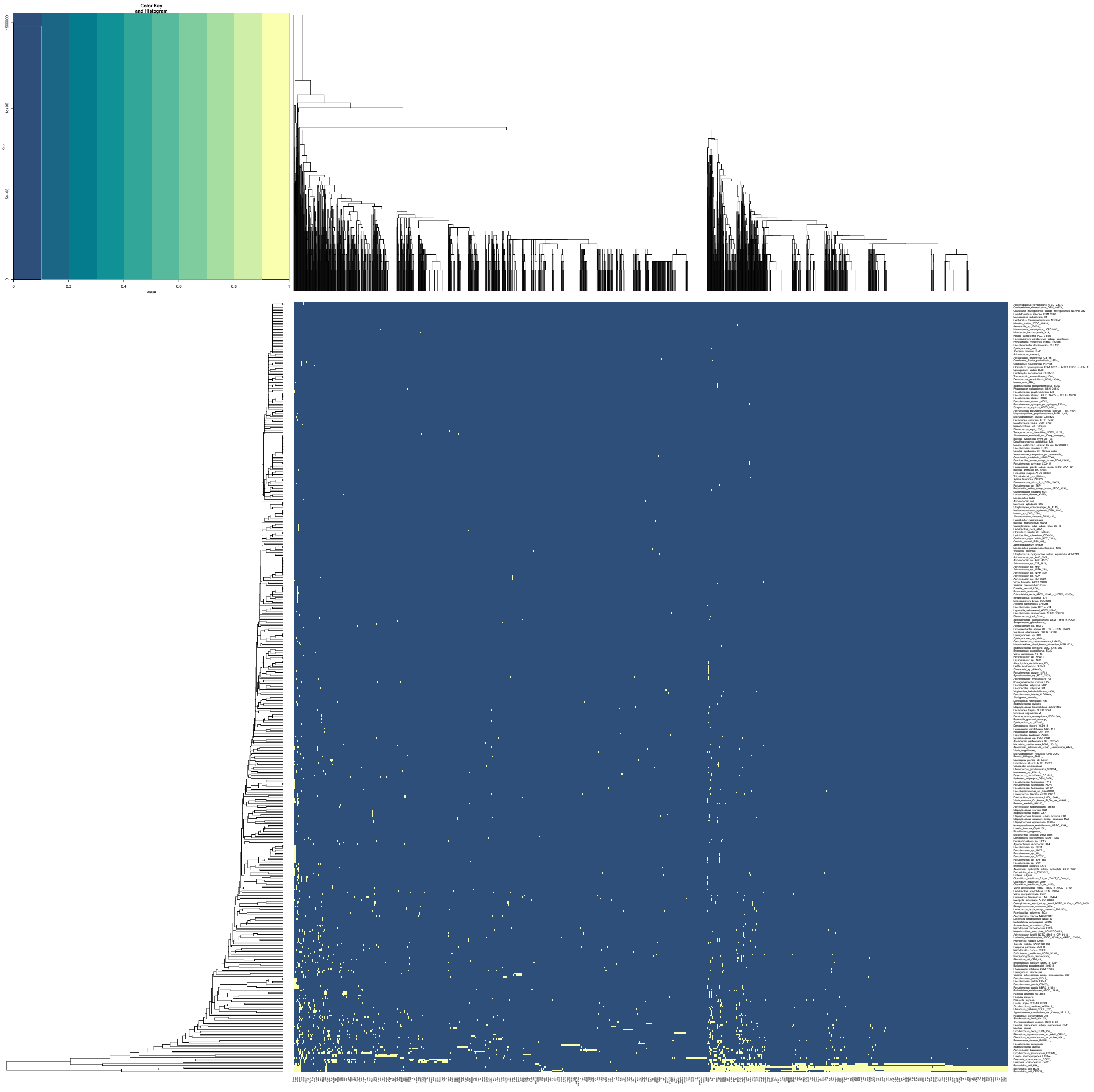


**Figure S2**. Heatmap visualising plasmid-encoded genes for 491 bacterial samples (y-axis) across 3,023 unique plasmid-encoded genes (x-axis). Dendrograms indicate the similarity across the samples (left) and genes (top). Both axes were sorted based on similarity. Genes are shown in yellow, and absence is show in blue. The yellow area of shared plasmid-encoded genes at the bottom right represents seven *E. coli*. See full figure on FigShare at https://doi.org/10.6084/m9.figshare.19525453.

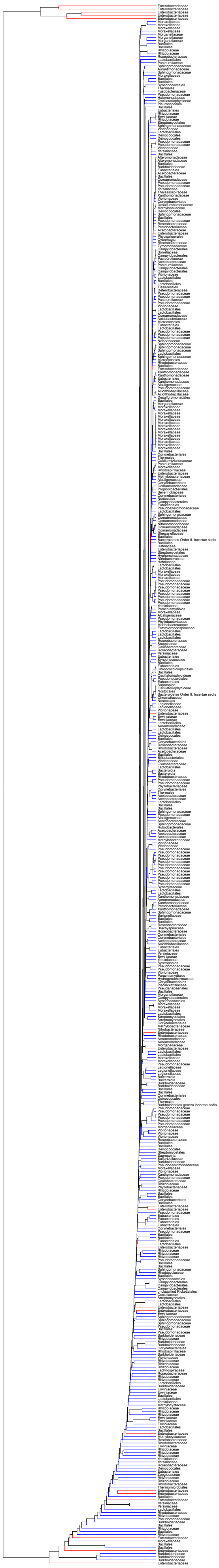

**Figure S3**. Dendrogram based on clustering of the 491 samples' taxonomic families based on their similarity across 3,023 plasmid-encoded genes indicating the classification of the different families. The *Enterobacteriaceae* are in red and non-*Enterobacteriaceae* are in blue.

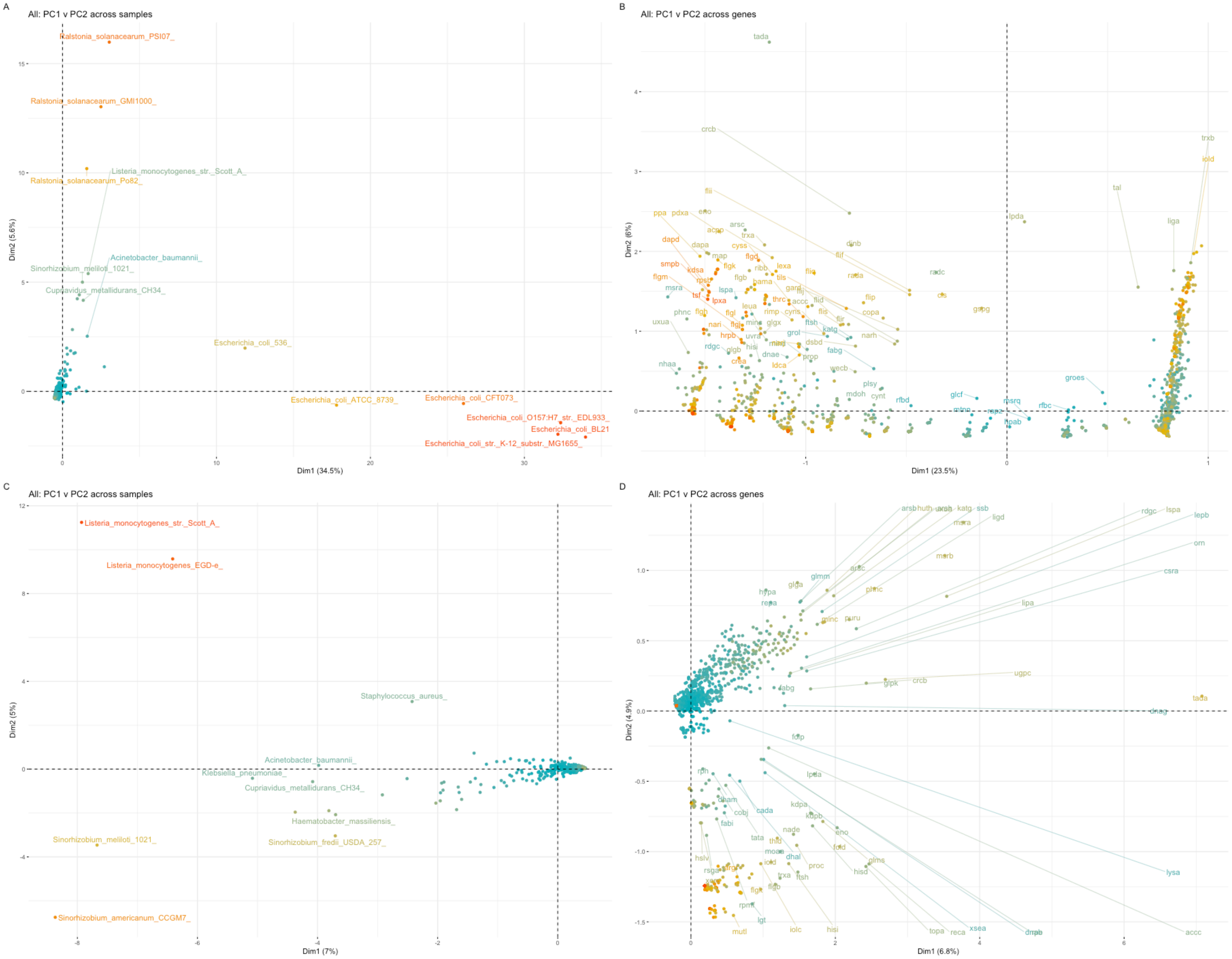


**Figure S4**. PCA showing the covariation of (A) the 497 samples based on their patterns across their 3,023 plasmid-encoded genes, (B) these plasmid-encoded genes based on their frequencies across the 497 samples, (C) 488 samples (excluding six *E. coli* and three *Ralstonia solanacearum* samples) based on their patterns across the plasmid-encoded genes, and (D) these plasmid-encoded genes based on their frequencies across the 488 samples. (C/D) Dim1 indicates PC1 (7.0% across samples, 6.8% across genes), and Dim2 indicates PC2 (5.0% across samples, 4.9% across genes). Lower PCs had much less of the total variation. The colours are cosmetic and are to differentiate different points

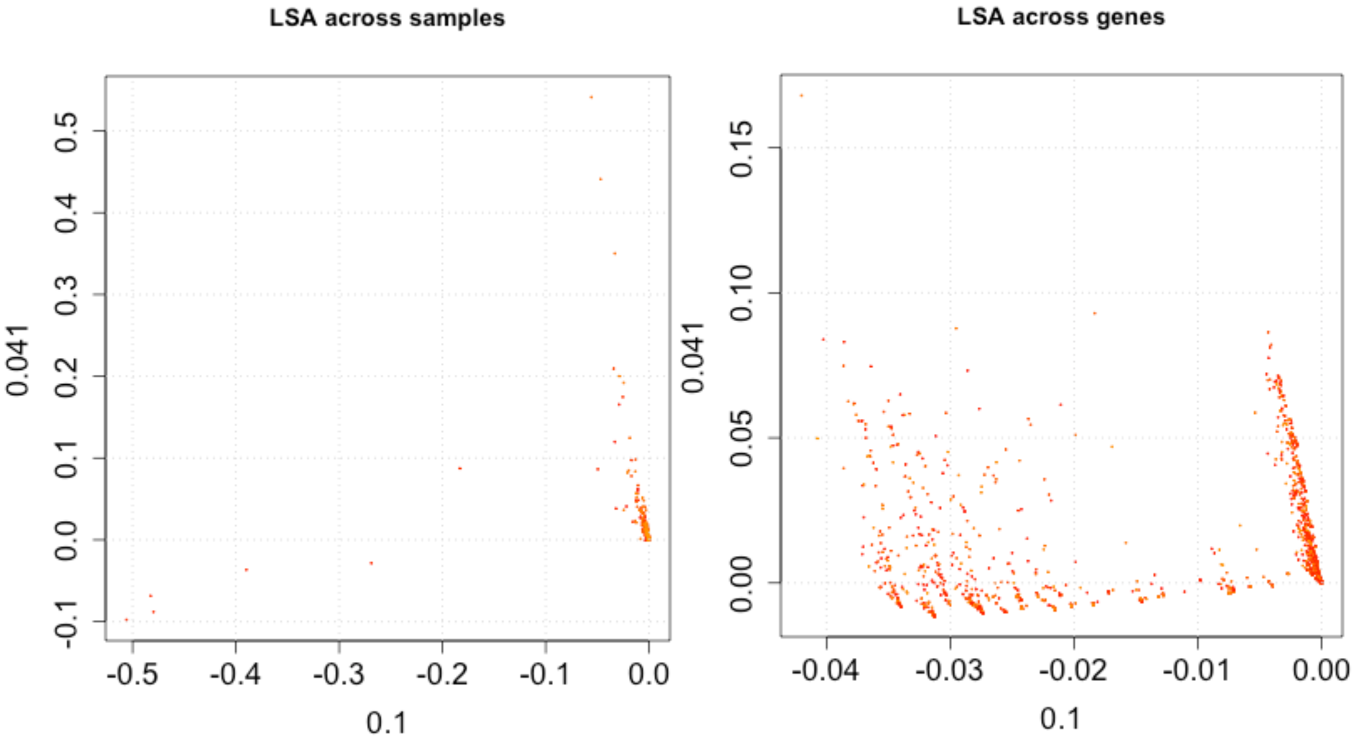


**Figure S6**. Latent semantic analysis (LSA) of 494 bacterial samples across 360 dimensions representing their 3,023 plasmid-encoded genes showing the variation across samples (left) and genes (right). The 1st dimension (x-axes) had 10.0% of variation, and the 2nd had 4.1% (y-axes). Left: The six *E. coli* samples are towards the bottom left and represent most of the variation across dimension 1; and the three *Ralstonia solanacearum* samples are towards the top right and represent most of the variation across dimension 2.

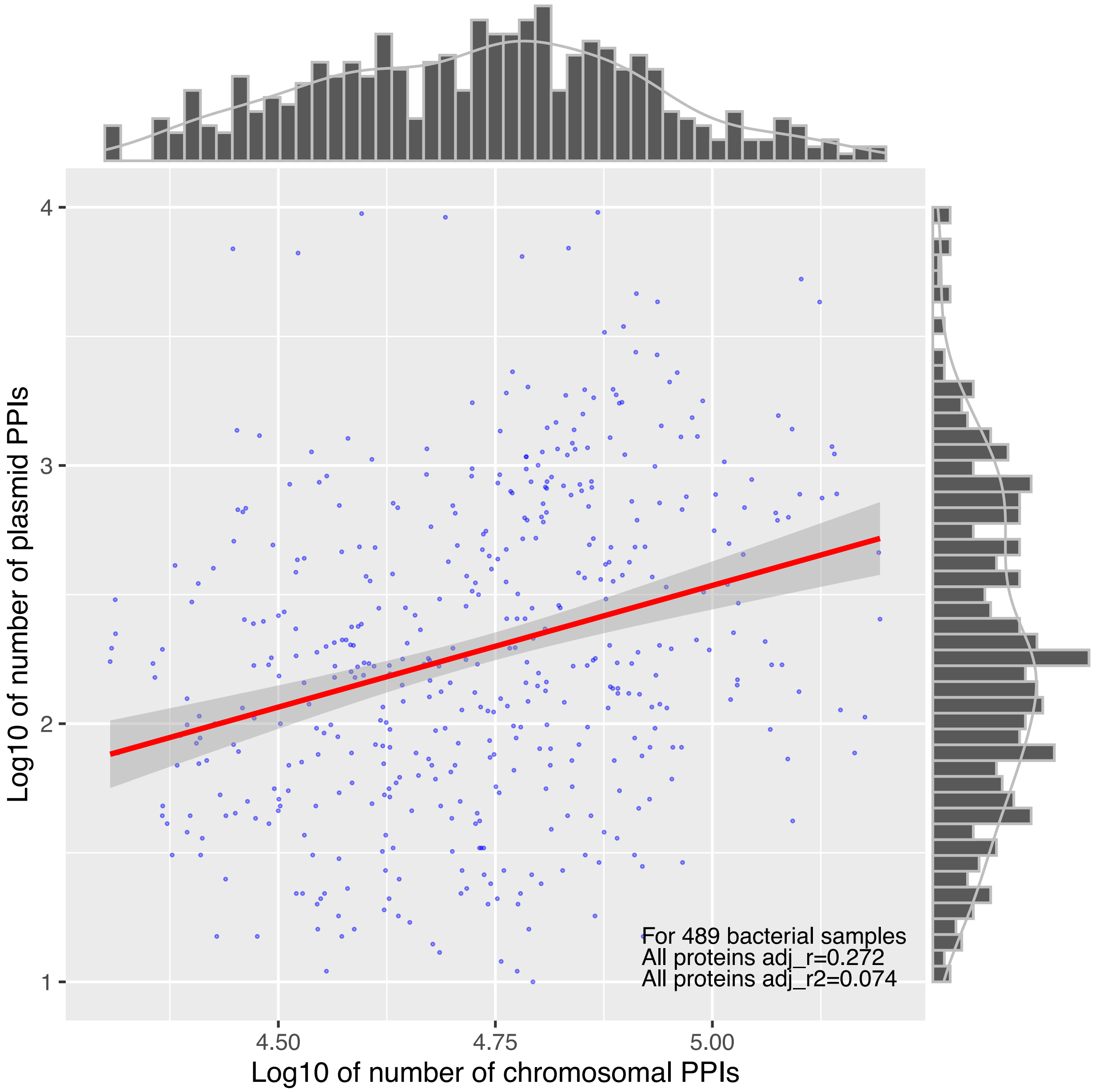


**Figure S6**. A positive correlation between the log10-scaled numbers of plasmid- (y-axis) and chromosome-related (x-axis) PPIs among 489 samples (purple points) with at least one plasmid-related PPI. The line of best fit (red) shows the linear association of the log10 of the number of plasmid- and chromosomal-related (x-axis) PPIs ($r=0.272$, $r^2=0.074$). The histograms indicate the marginal densities for each axis.

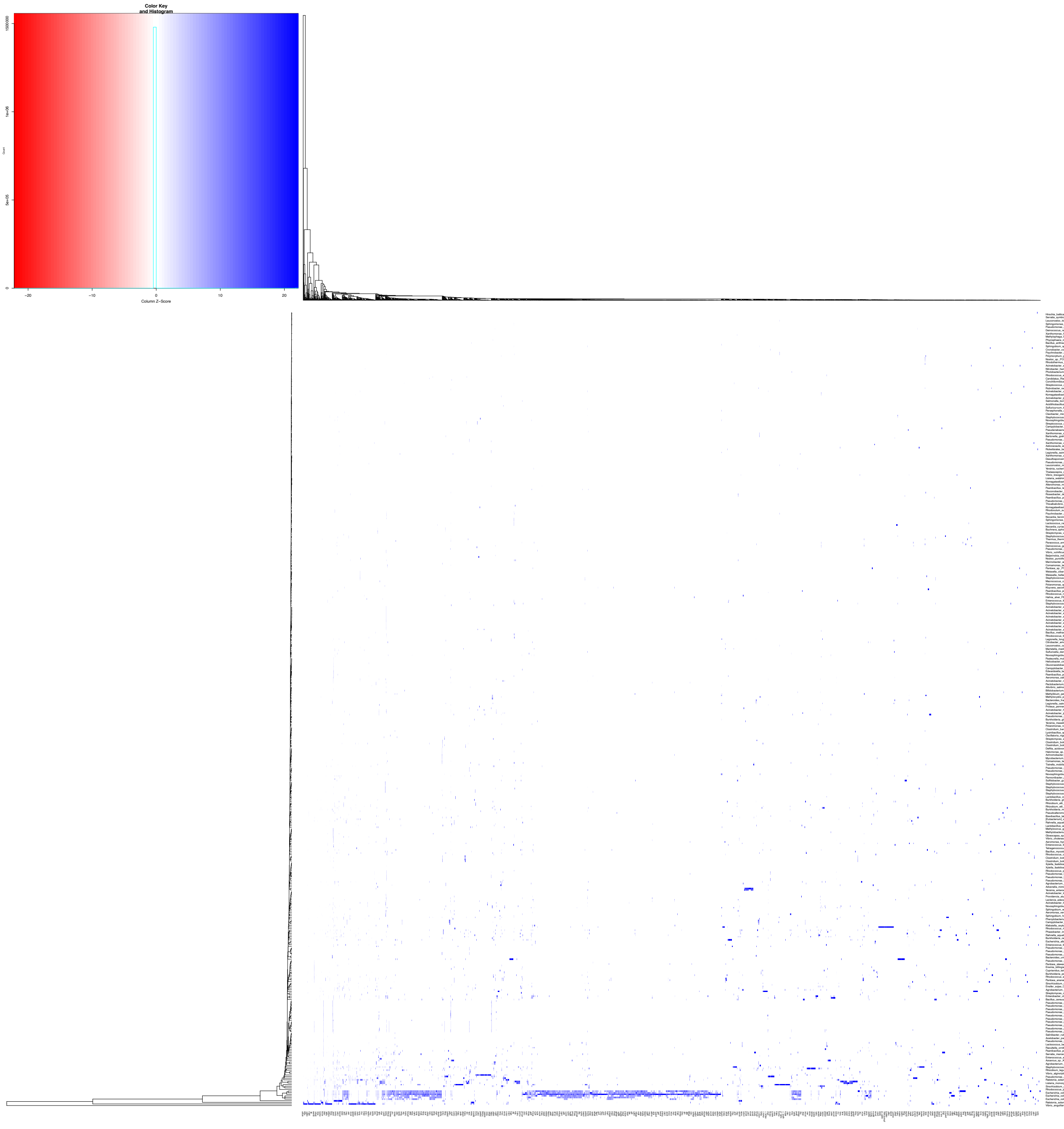


**Figure S7**. Heatmap visualising plasmid interactomes for 489 bacterial samples with at least one plasmid-related PPI (y-axis) across their 3,023 unique plasmid-derived genes' PPI rates (x-axis). The values are normalised by column. See full figure on FigShare at https://doi.org/10.6084/m9.figshare.19525465 and data at https://doi.org/10.6084/m9.figshare.19525471.

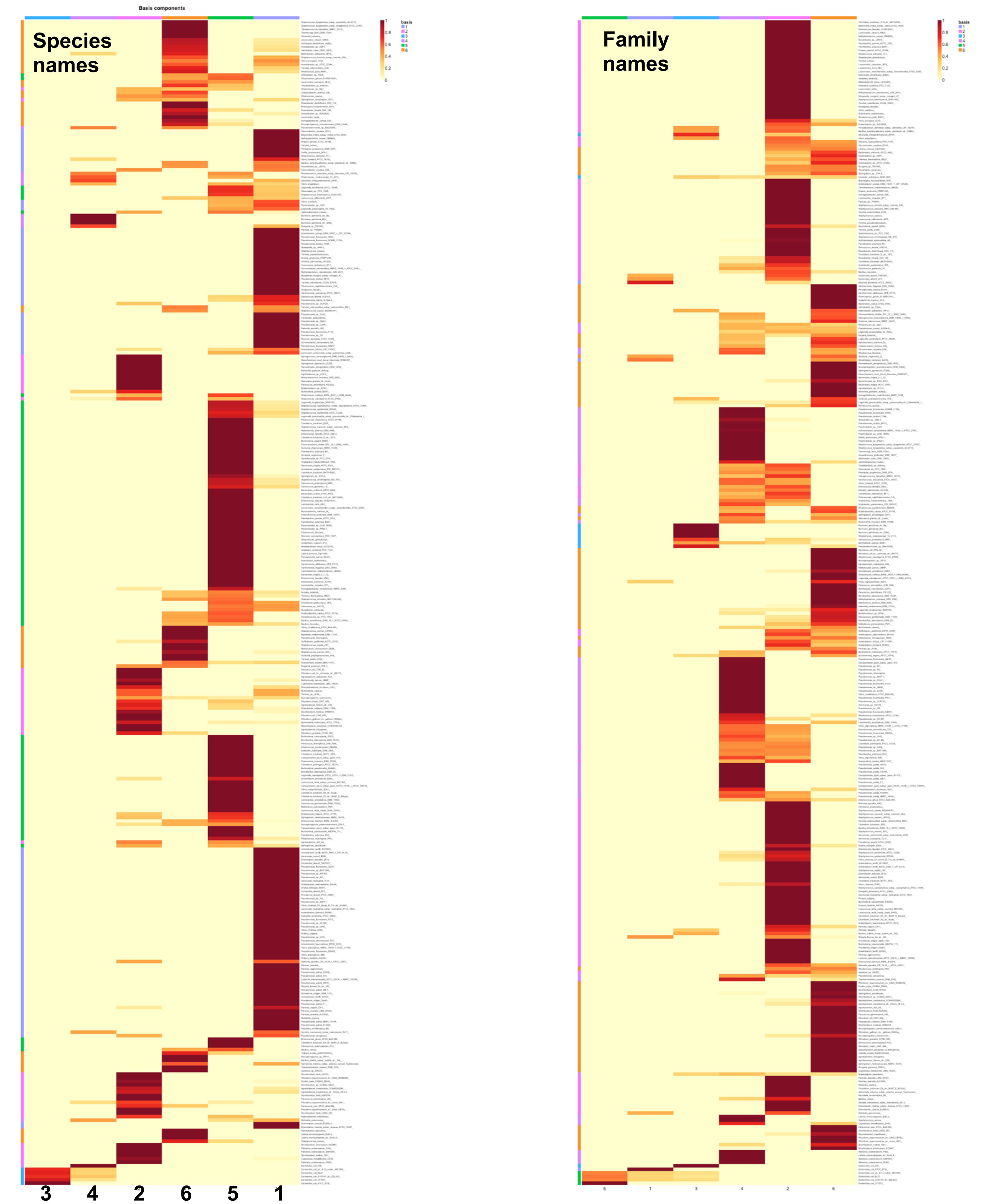


**Figure S8**. Heatmaps of the basis coefficient matrices, reflecting the probability of allocation of the 331 samples (y-axes) to seven different groups (x-axes) based on their presence-absence profiles in 1,733 plasmid-encoded genes using NMF using their species names (left) and family names (right, from a different NMF run). The legend below shows the probability of allocation (as per the legend: high in red, low in yellow) to the different groups ("basis"). Left: 160, 71, 5, 21 136 and 71 samples had high probabilities of being allocated to each respective rank. Rank 5 (right) is equivalent to the initial rank 3 with the five *E. coli* (left). See full left figure on FigShare at https://doi.org/10.6084/m9.figshare.19525492, data at https://doi.org/10.6084/m9.figshare.19525480.

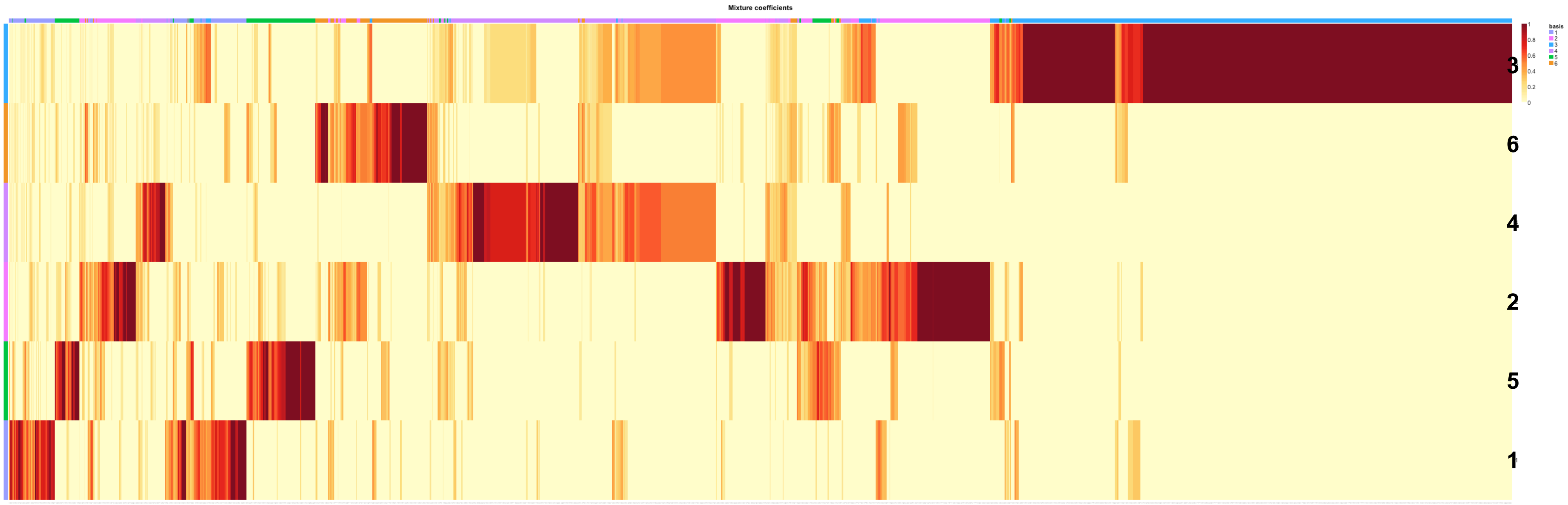


**Figure S9**. Heatmap of the mixture coefficient matrices, reflecting the probability of allocation of 1,733 plasmid-encoded genes (x-axis) to six different groups (y-axis) based on their presence or absence in 331 samples using NMF. The legend below shows the probability of allocation (as per the legend: high in red, low in yellow) to the different groups ("basis"). 286, 584, 1,065 (corresponding to the five *E. coli*), 482, 324 and 416 plasmid-encoded genes had high probabilities of being allocated to each respective rank. See full figure on FigShare at https://doi.org/10.6084/m9.figshare.19525591 and data at https://doi.org/10.6084/m9.figshare.19525597

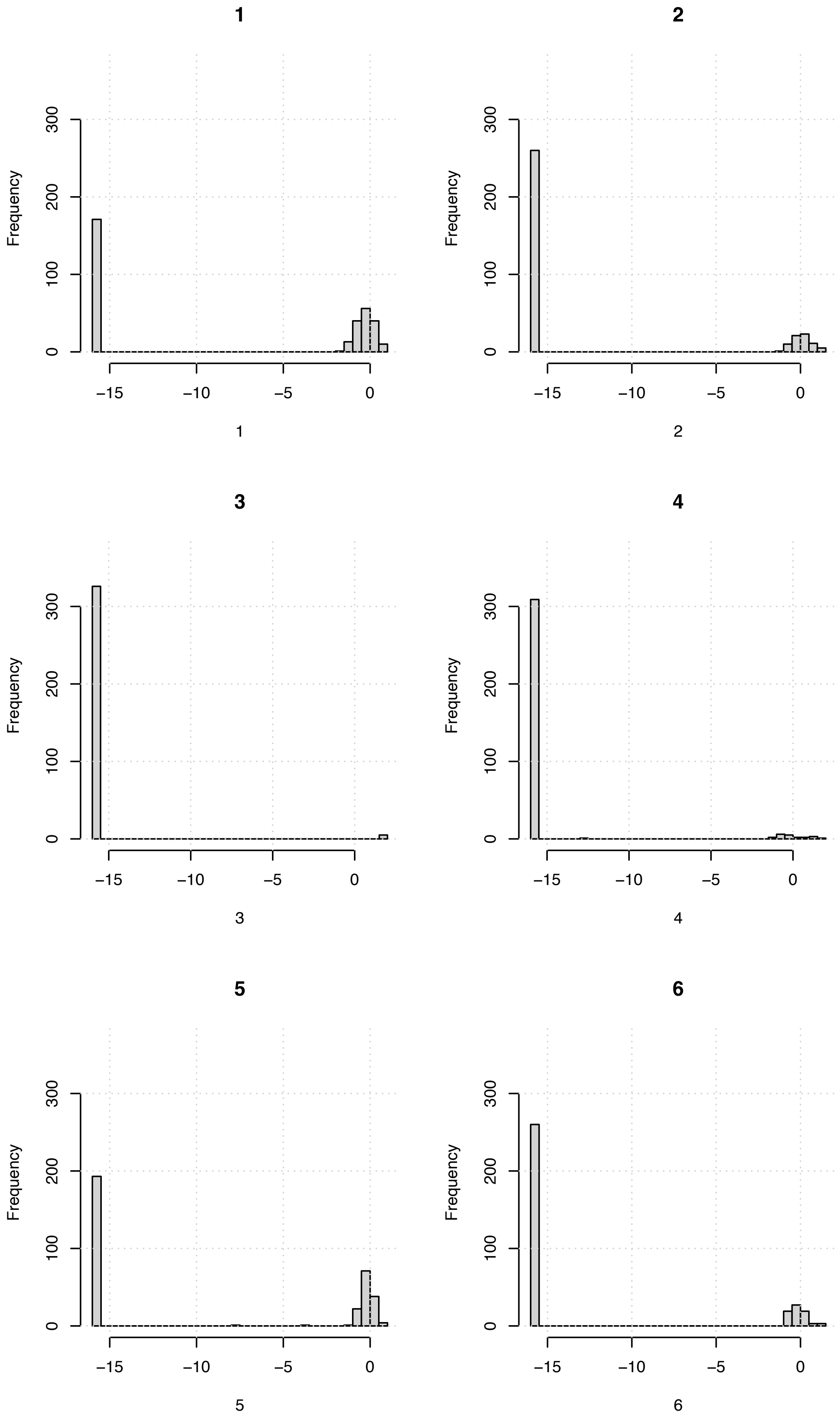


**Figure S10**. The frequency (y-axis) of the log10-scaled probabilities of allocating (x-axis) the 331 samples to each of seven ranks from NMF based on their presence-absence rates in 1,733 plasmid-encoded genes. 160, 71, 5 (corresponding to the five *E. coli*), 21 136 and 71 samples had high probabilities of being allocated to each respective rank.

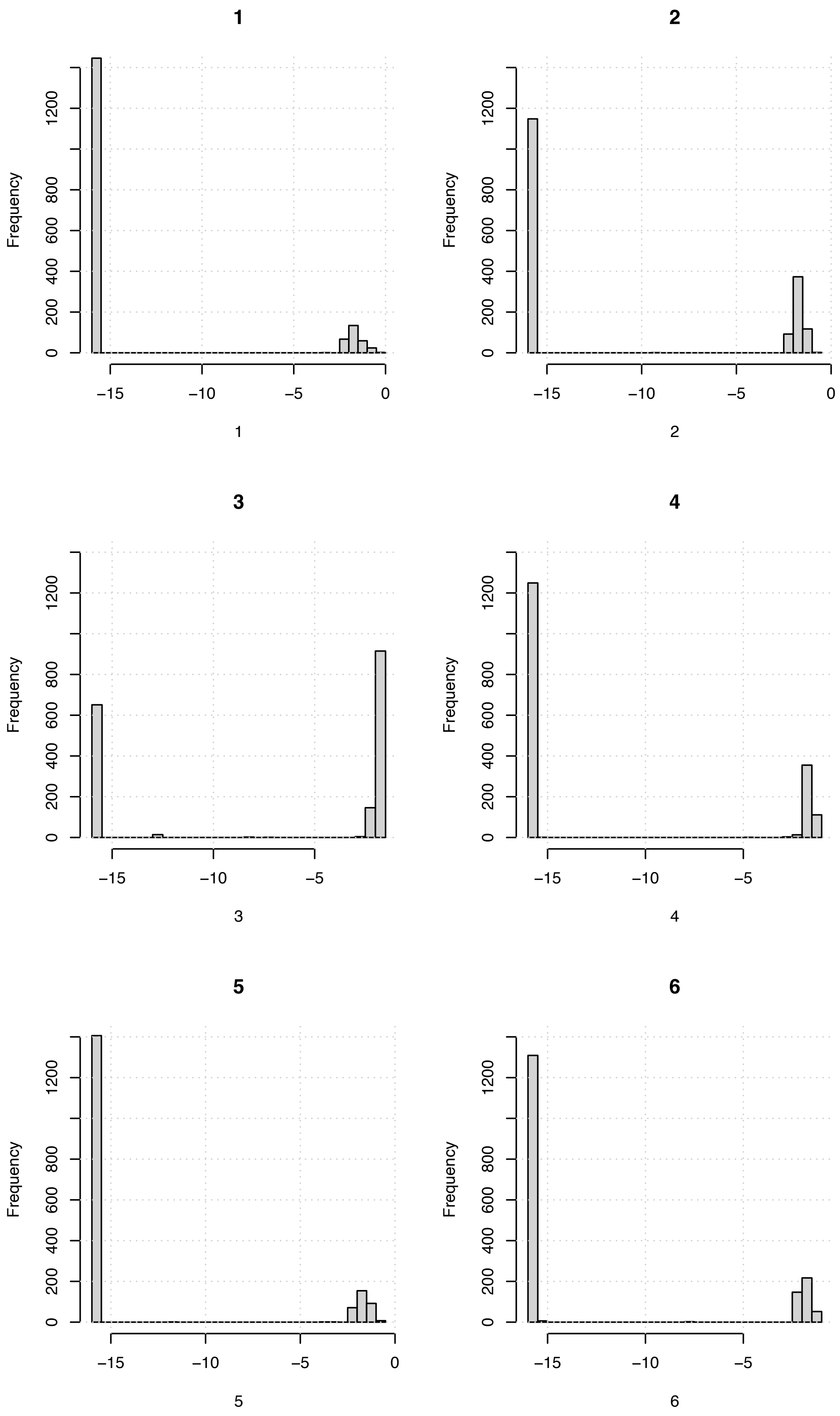


**Figure S11**. The frequency (y-axis) of the log10-scaled probabilities of allocating (x-axis) the 1,733 plasmid-encoded genes to each of six ranks from NMF based on their presence-absence rates in 331 samples. 286, 584, 1,065 (corresponding to the five *E. coli*), 482, 324 and 416 plasmid-encoded genes had high probabilities of being allocated to each respective rank.

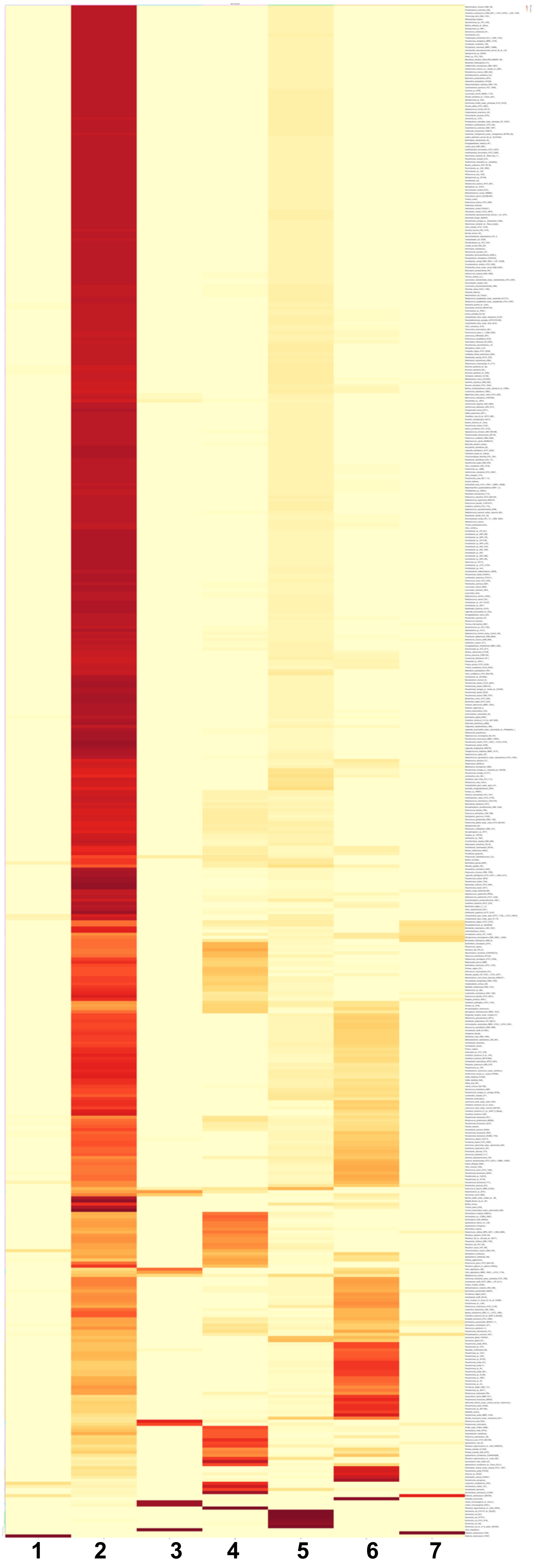


**Figure S12**. Heatmap of the basis coefficient matrices, reflecting the probability of allocation of the 481 samples (y-axis) to seven different groups (x-axis) based on their plasmid gene PPI rates in 2,363 plasmid-encoded genes using NMF. The legend below shows the probability of allocation (as per the legend: high in red, low in yellow) to the different groups ("basis"). The six *E. coli* samples were allocated to rank 5. See full figure on FigShare at https://doi.org/10.6084/m9.figshare.19611276 and data at https://doi.org/10.6084/m9.figshare.19611279.

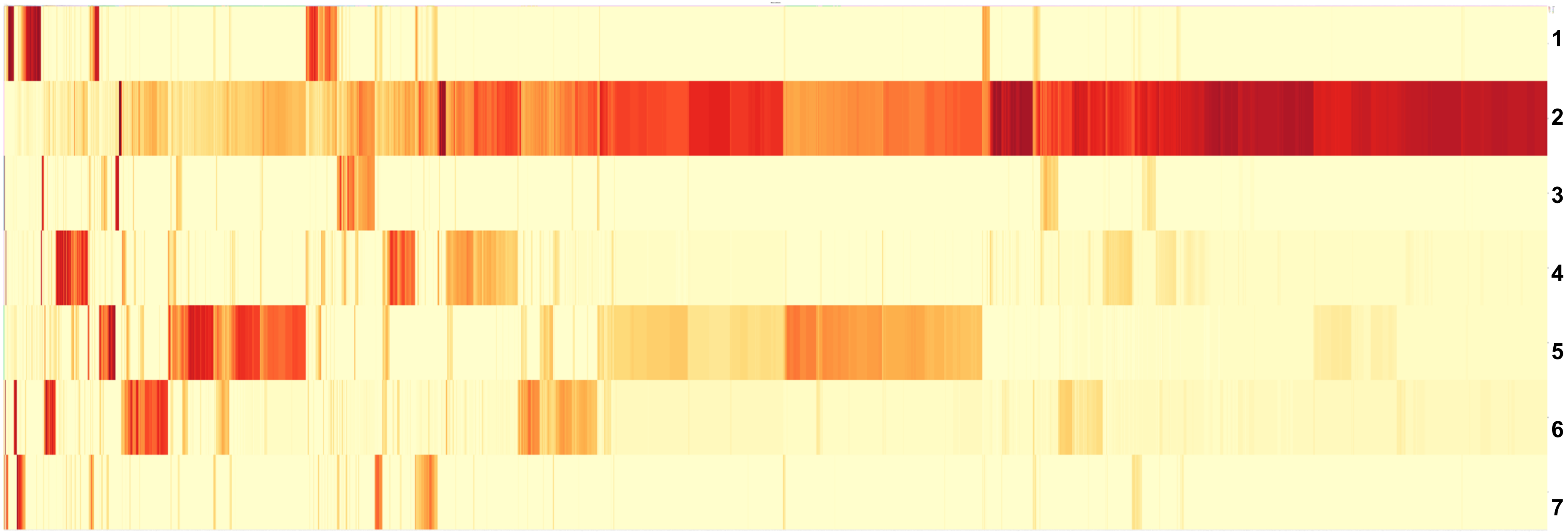


**Figure S13**. Heatmap of the mixture coefficient matrices, reflecting the probability of allocation of 2,363 plasmid-encoded genes (x-axis) to seven different groups (y-axis) based on their PPI rates in 481 samples using NMF. The legend below shows the probability of allocation (as per the legend: high in red, low in yellow) to the different groups ("basis"). Rank 5 corresponded to 238 protein's PPI rates. The six *E. coli* samples' genes were allocated to rank 5. See full figure on FigShare at https://doi.org/10.6084/m9.figshare.19611282 and data at https://doi.org/10.6084/m9.figshare.196'1285 .

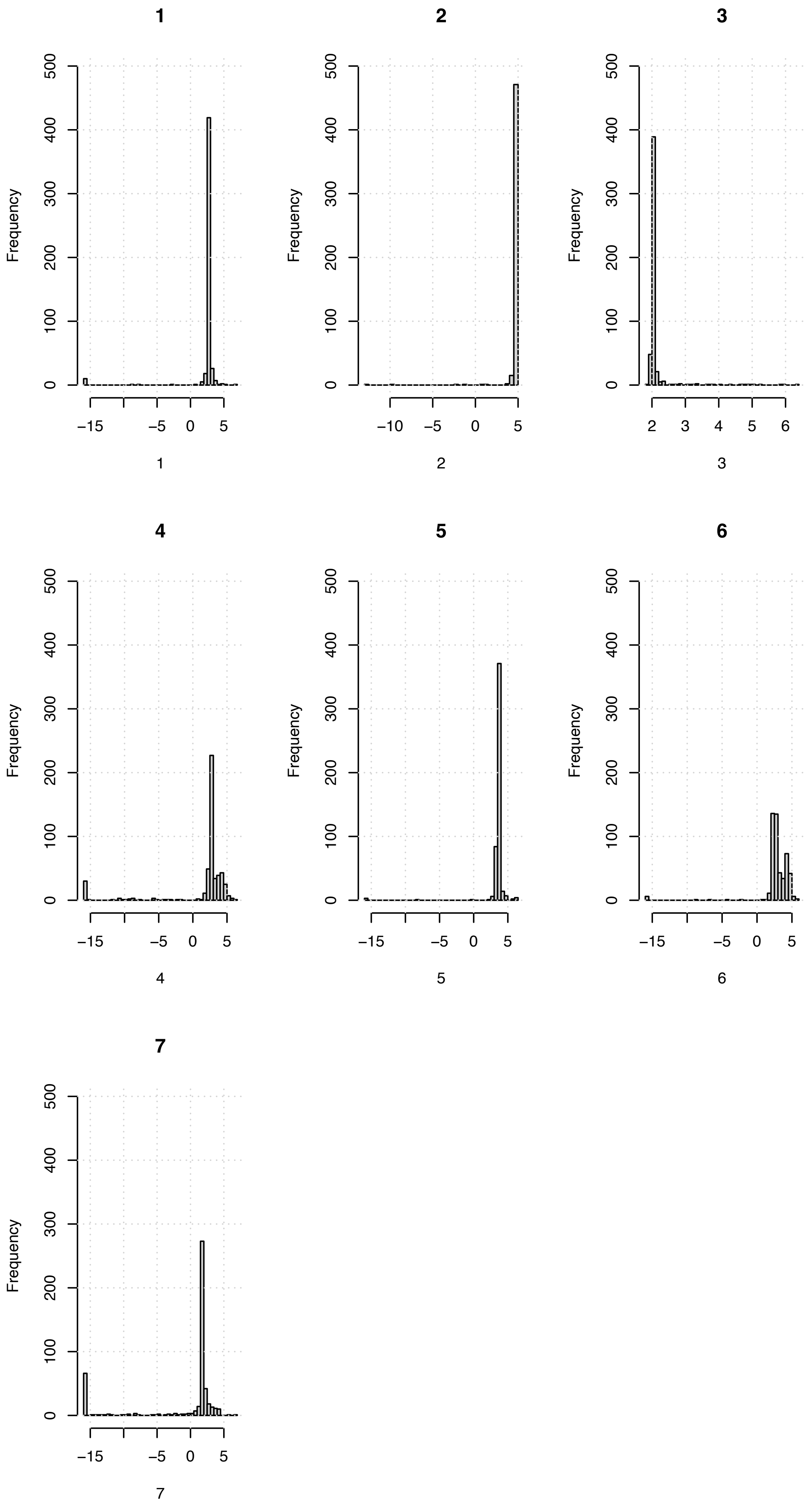


**Figure S13**. The frequency (y-axis) of the log10-scaled probabilities of allocating (x-axis) the 481 samples to each of seven ranks from NMF based on their PPI rates in 2,363 plasmid-encoded genes. The six *E. coli* samples were allocated to rank 5.

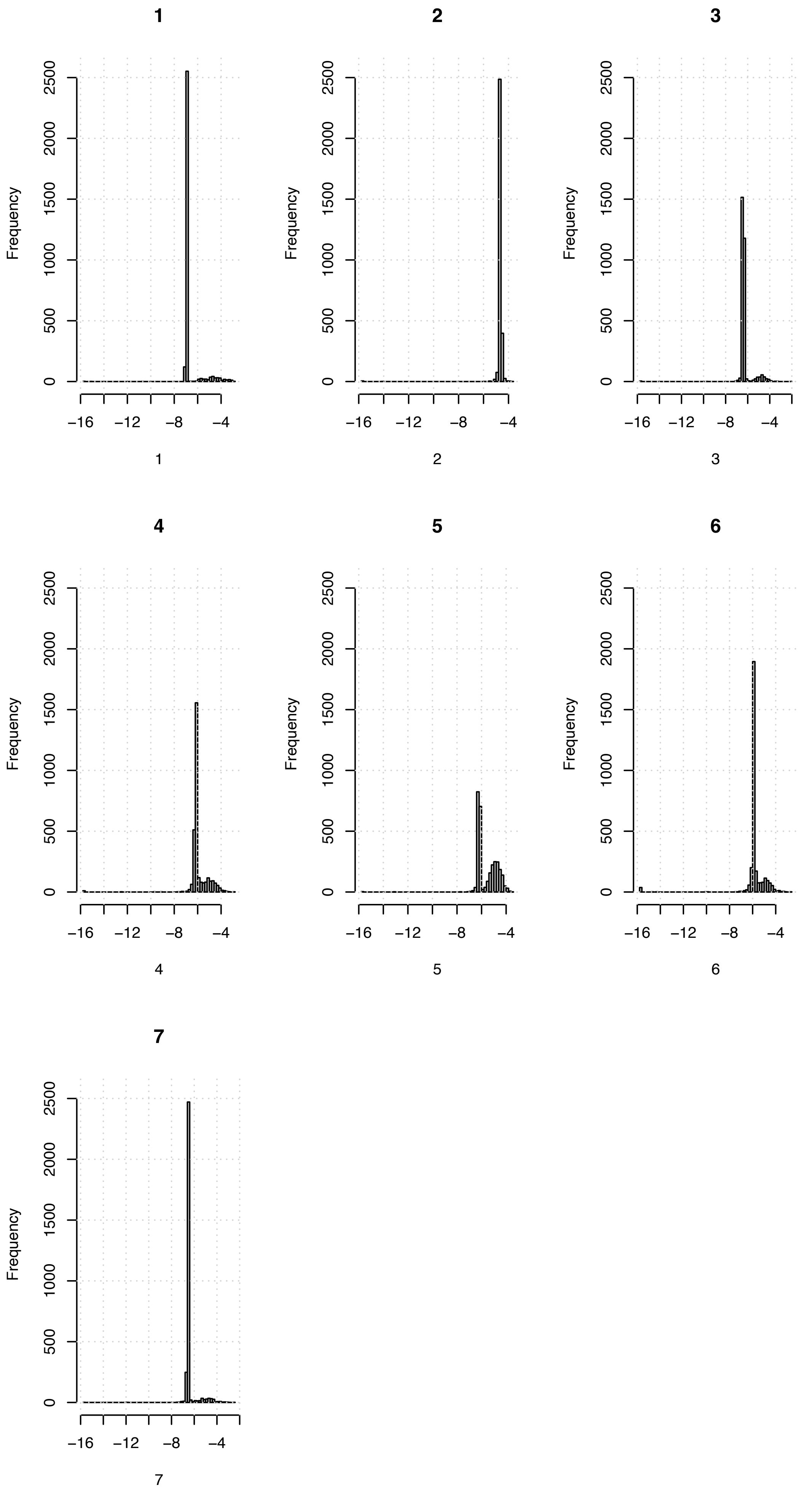


**Figure S14**. The frequency (y-axis) of the log10-scaled probabilities of allocating (x-axis) the 2,363 plasmid-encoded genes to each of seven ranks from NMF based on their PPI rates in 481 samples. The six *E. coli* samples' proteins' PPI rates were

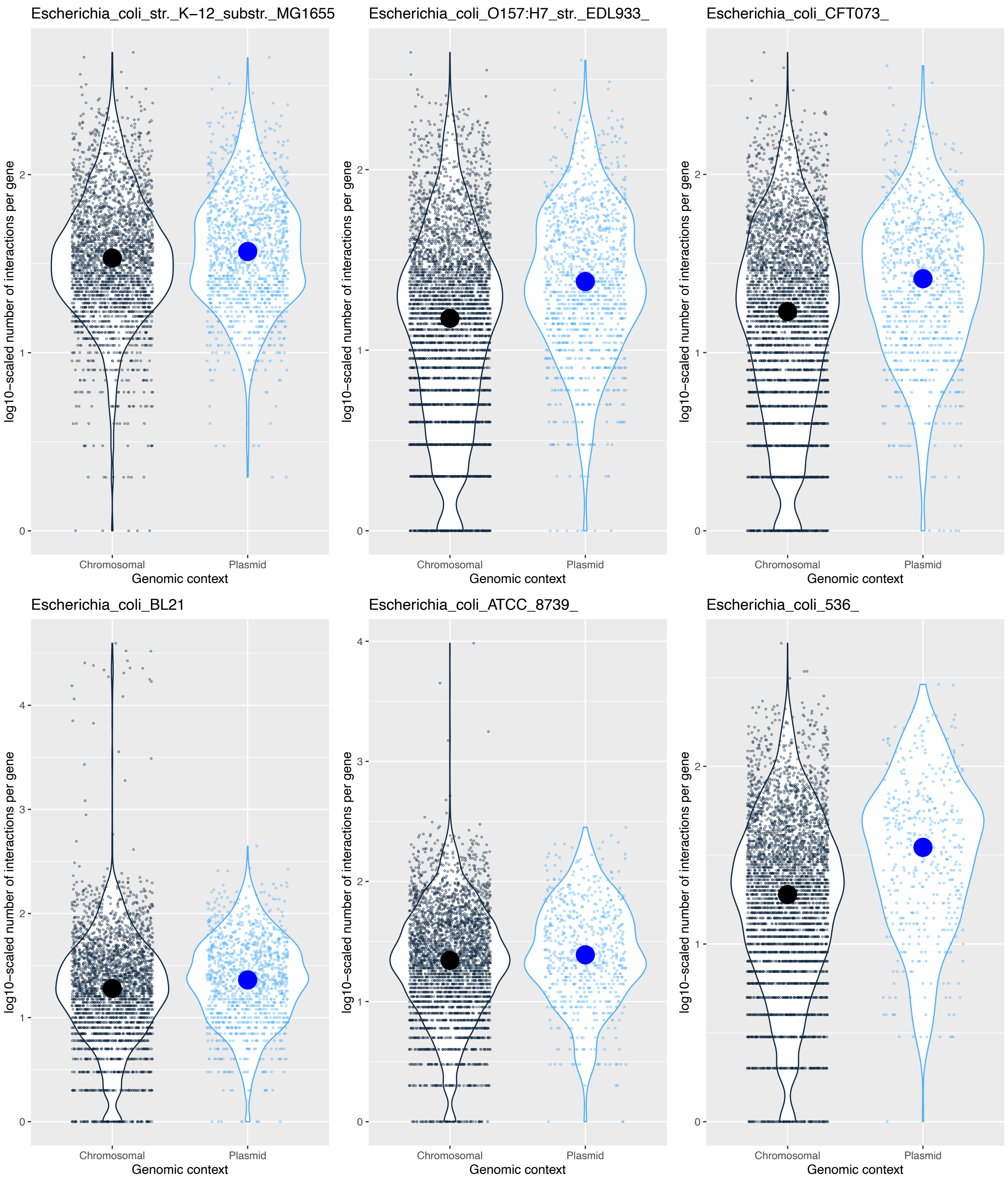


**Figure S15**. The log10-scaled numbers of chromosomal (black) and plasmid (blue) PPIs per gene for six *E. coli*: K-12 substrain MG1665 (top left), O157:H7 substrain EDL933 (top middle), CFT073 (top right), BL21 (bottom left), ATCC8793 (bottom middle), and 536 (bottom right). The data for these and the full set of chromosomal and plasmid interactions per gene for the samples are available as 4,430 CSV files and 344 PDF images (images were generated if there was sufficient PPI data to plot) on FigShare at doi: https://doi.org/10.6084/m9.figshare.19576408. The points show the observed values on top of the distribution shapes and the medians shown by large circles.

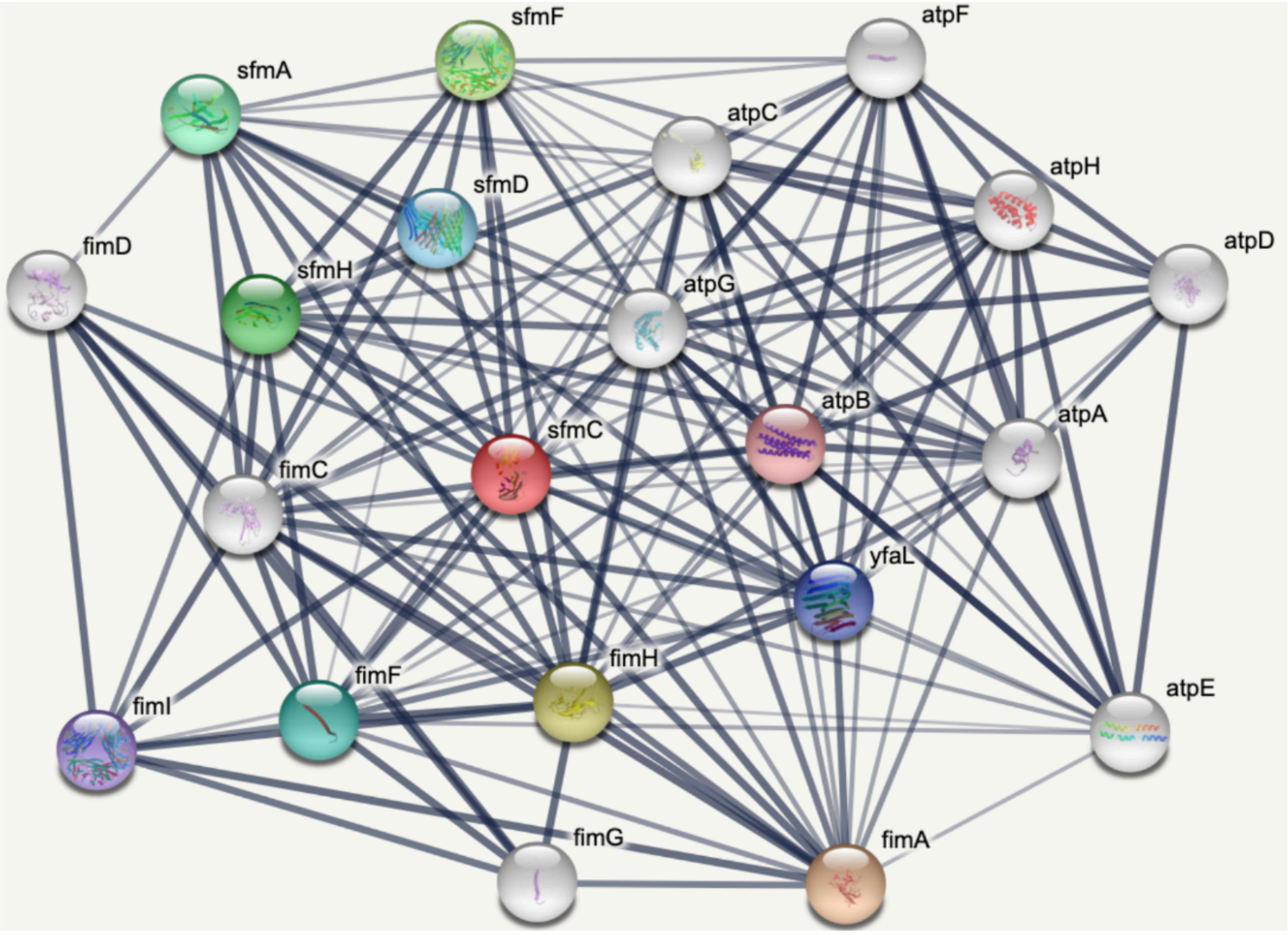


**Figure S16**. An *E. coli* K12 MG1655 (String ID 511145) PPI network from the StringDB website of 21 proteins in Table S4 centred on SfmC (a putative periplasmic pilus chaperone that is part of the *sfmACDHF* fimbrial operon) with 816 PPIs in total.

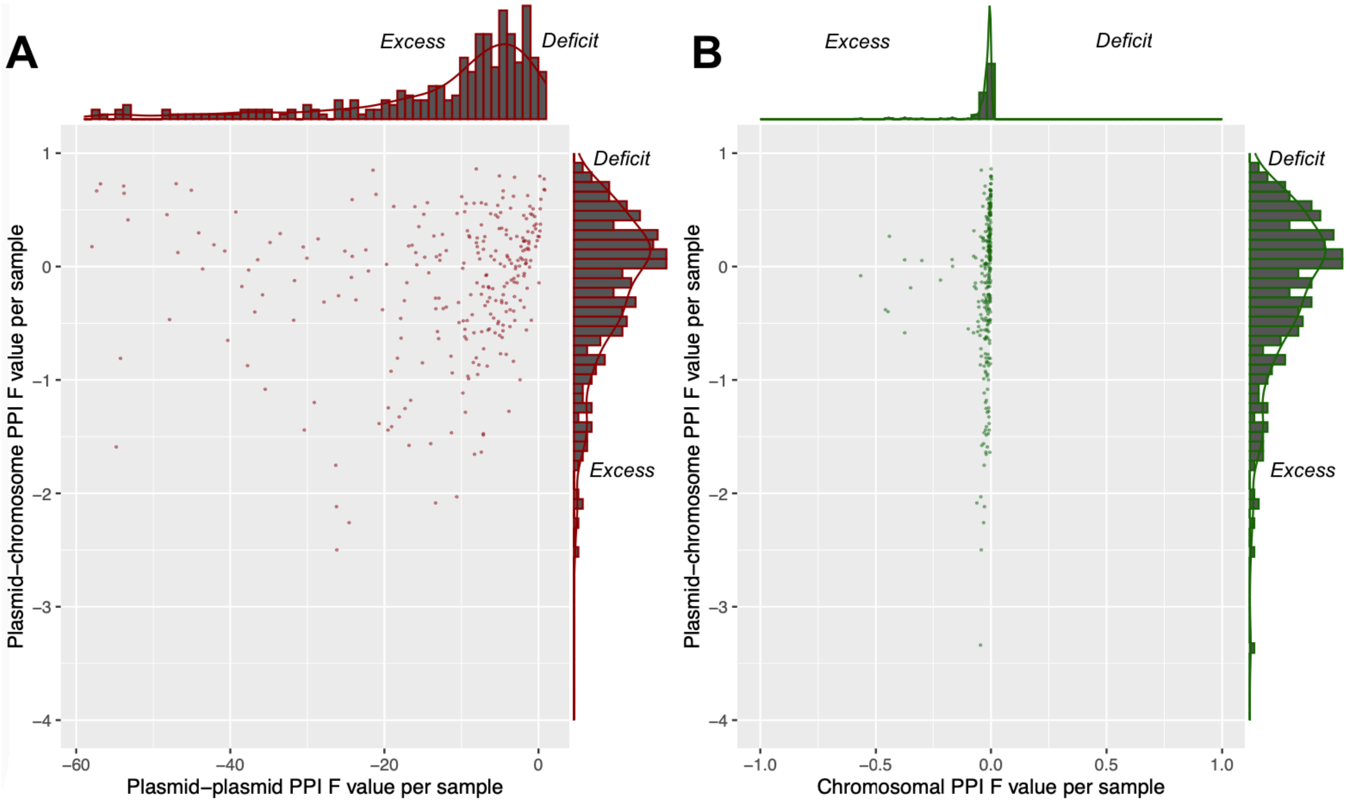


**Figure S18**. The F values for plasmid-plasmid PPIs per sample (x-axis) versus (A) the F values for plasmid-chromosome PPIs per sample (y-axis) (red points) and (B) the F values for chromosomal PPIs per sample (y-axis) (green points) in 283 samples with at least one PPI involving only plasmid-encoded proteins. F > 0 indicates a deficit of the PPI type per sample, F < 1 indicates an excess of that PPI type per sample. Plasmid-plasmid PPIs show much higher rates of excess number compared to (A) plasmid-chromosome and (B) chromosomal PPIs. The histograms indicate the marginal densities per axis.

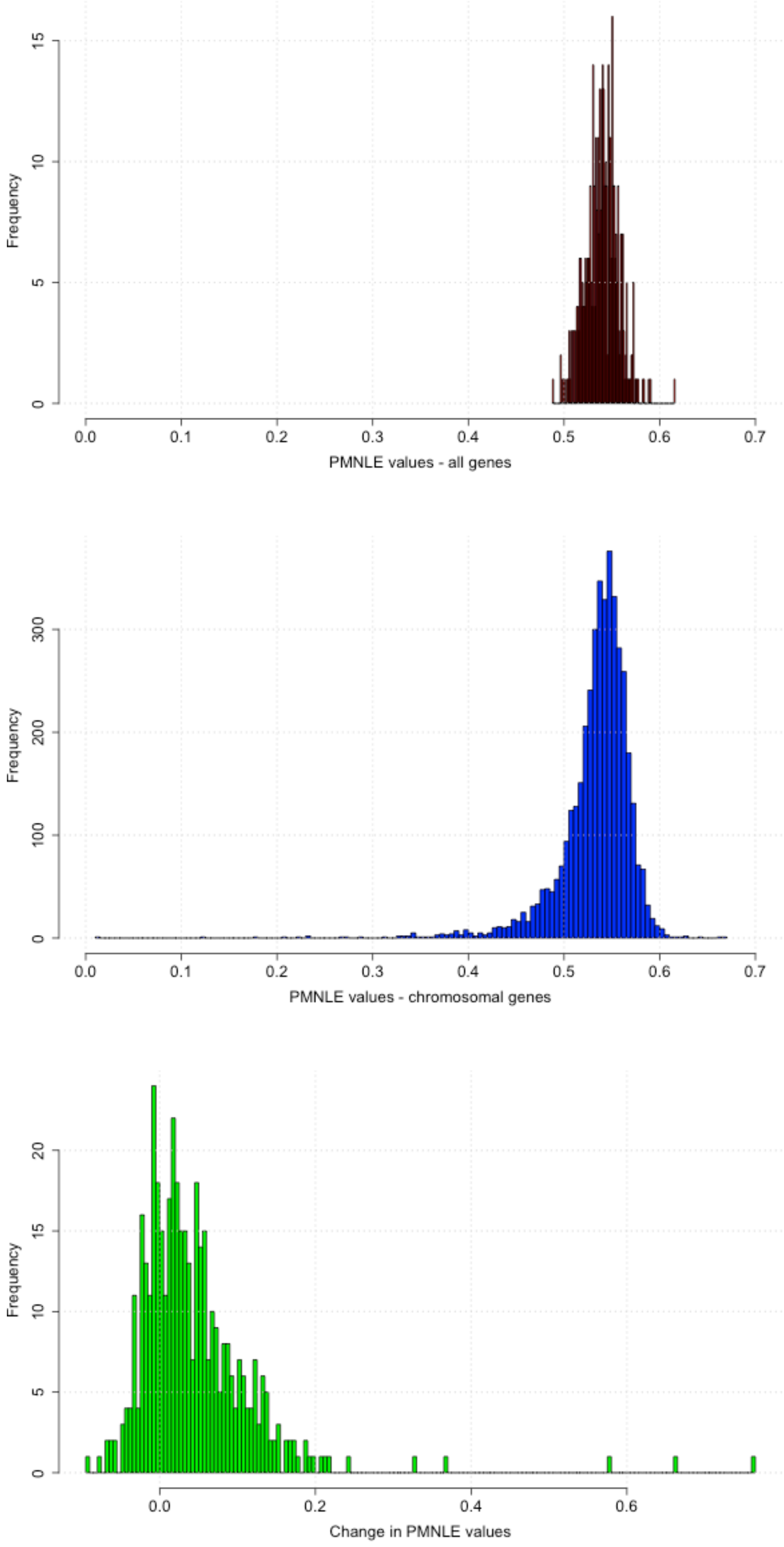


**Figure S19**. The distribution of aPMNLE values (x-axis) for all genes in the 489 samples with plasmid-related PPIs (top, red), aPMNLE values for chromosomal genes in 4,363 samples (middle, blue) and the scaled difference between all and chromosomal genes in the 489 samples with plasmid-related PPIs (bottom, green). Note that the y-axes scales differ.

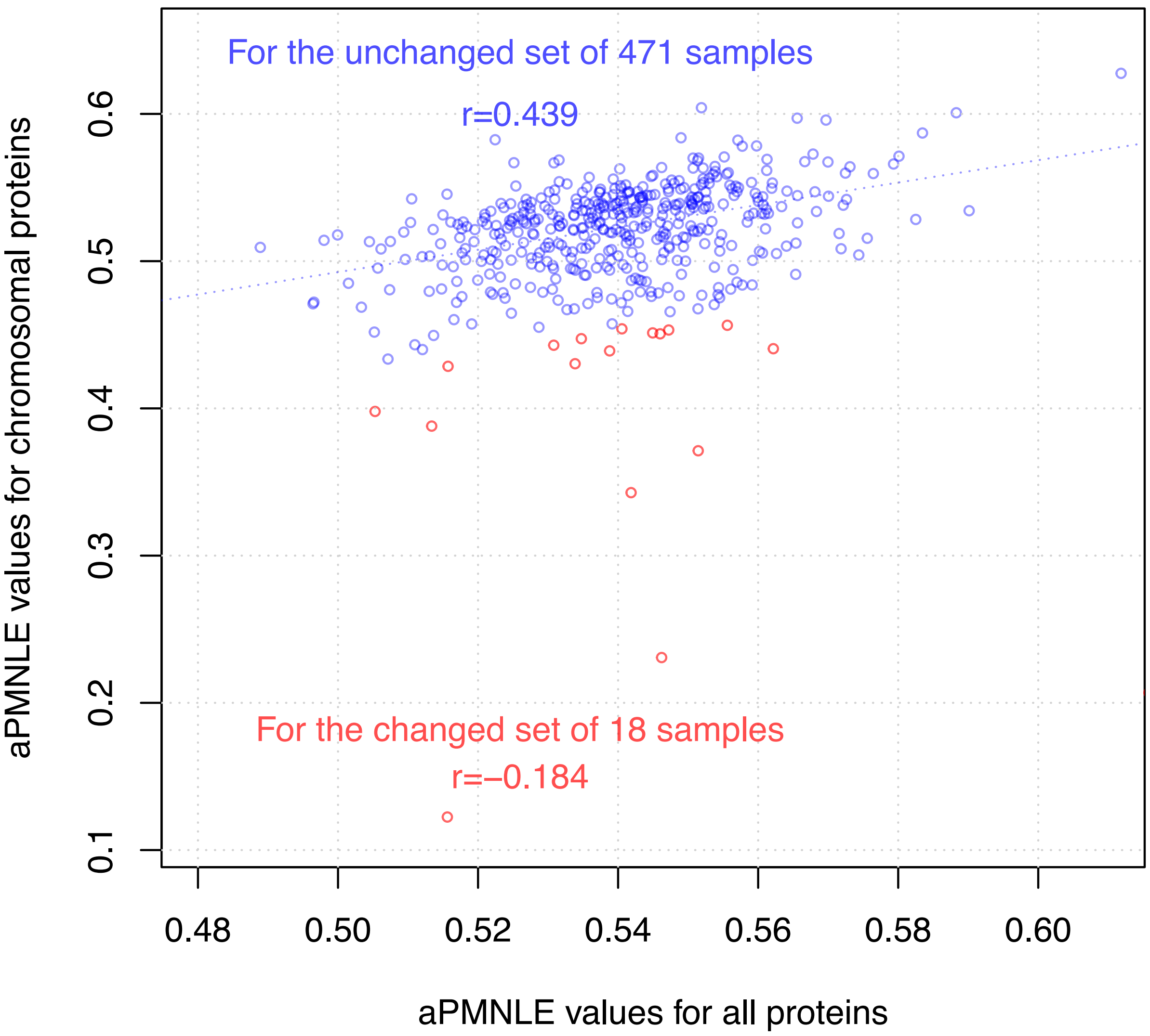


**Figure S20**. The association between the aPMNLE values for all (chromosomal and plasmid) proteins (x-axis) compared to those for chromosomal proteins (y-axis) for the group with no significant difference in these aPMNLE values ("unchanged", n=471 samples, blue) and the group where there was a much lower chromosomal aPMNLE value ("changed", n=18, red). The correlation between these aPMNLE metrics was higher for the main group of 471 samples (r=0.44, blue dashed line of best fit) compared to the group with the change in aPMNLE value (r=-0.18). The group of 18 had much lower chromosomal aPMNLE values when compared to the other 471 samples.

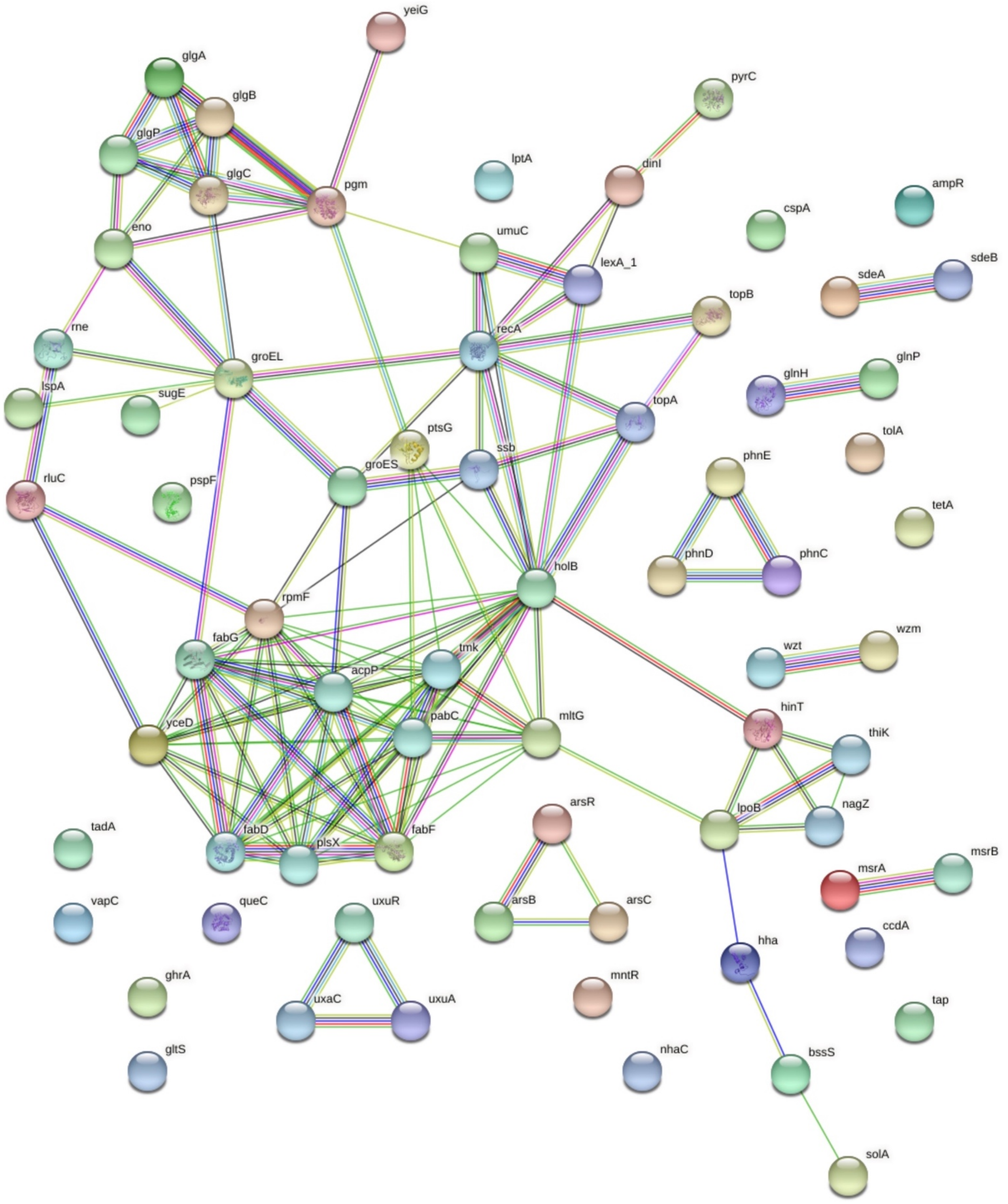


**Figure S21**. A PPI network of the 69 unique plasmid-encoded proteins in *Serratia marcescens* subsp marcescens Db11 that had 92 plasmid-restricted PPIs with one another. Chromosomal proteins are not shown here. This illustrates the extent to which plasmid-related proteins have PPIs less with one another, and more with chromosomal proteins.

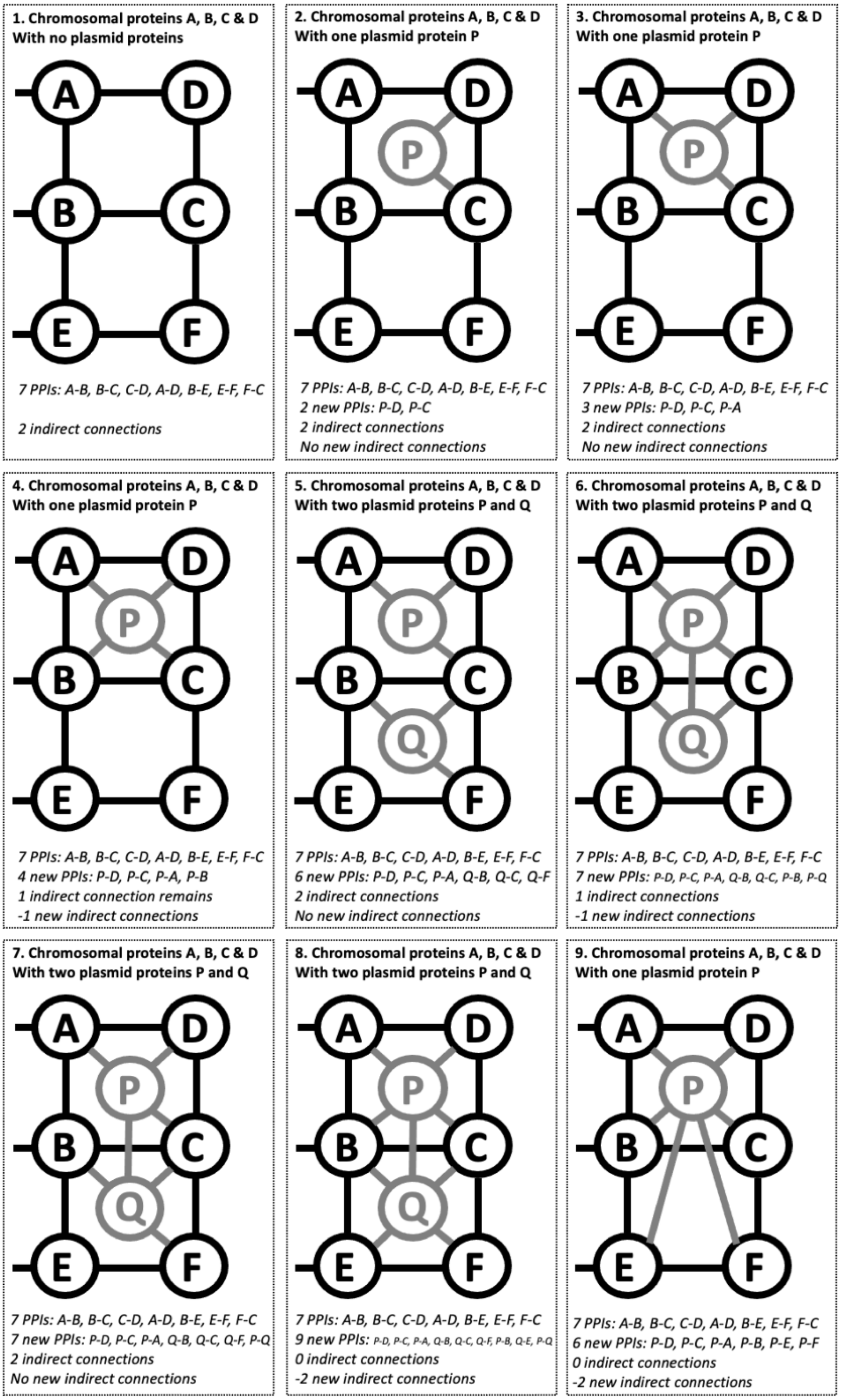


**Figure S22**. A model showing how PPI rates can increase but indirect connection numbers can fall simultaneously when plasmid-encoded proteins are added to a PPI network. (1) If four chromosomal proteins (A, B, C, D) form part of a PPI network such that they share seven PPIs between A-B, B-C, C-D, D-A, B-E, E-F and F-C, then they create two indirect connections (A-B-C-D and B-E-F-C). (2) If one plasmid-encoded protein P is added to this network such that it creates two new PPIs with D and C, it creates no new indirect connections. (3) If P also has a 3rd PPI with A, no change in the numbers of indirect connections is observed. (4) If P now has a 4th PPI with D, one indirect connection is lost because A, B, C, D and P are now connected directly by PPIs, but still creating four new PPIs (B-E-F-C remains as an indirect connection). (5) If we start with (3) and add a 2nd plasmid protein Q that has PPIs with B, C and Q, we have six new PPIs and no change in the indirect connections. (6) If we add PPIs between P-B and P-Q and lose one PPI at Q-F, we lost one indirect connection but created seven new PPIs (B-E-F-C remains as an indirect connection). (7) If we start with (5) and connect P-Q only, the numbers of indirect connections do not change. (8) If we extended (6) by connecting Q-E and Q-F, we have created nine new PPIs and lose both existing indirect connections. (9) If we revert to P only that has PPIs with all six chromosomal proteins, then we have six new PPIs and lose both indirect connections.

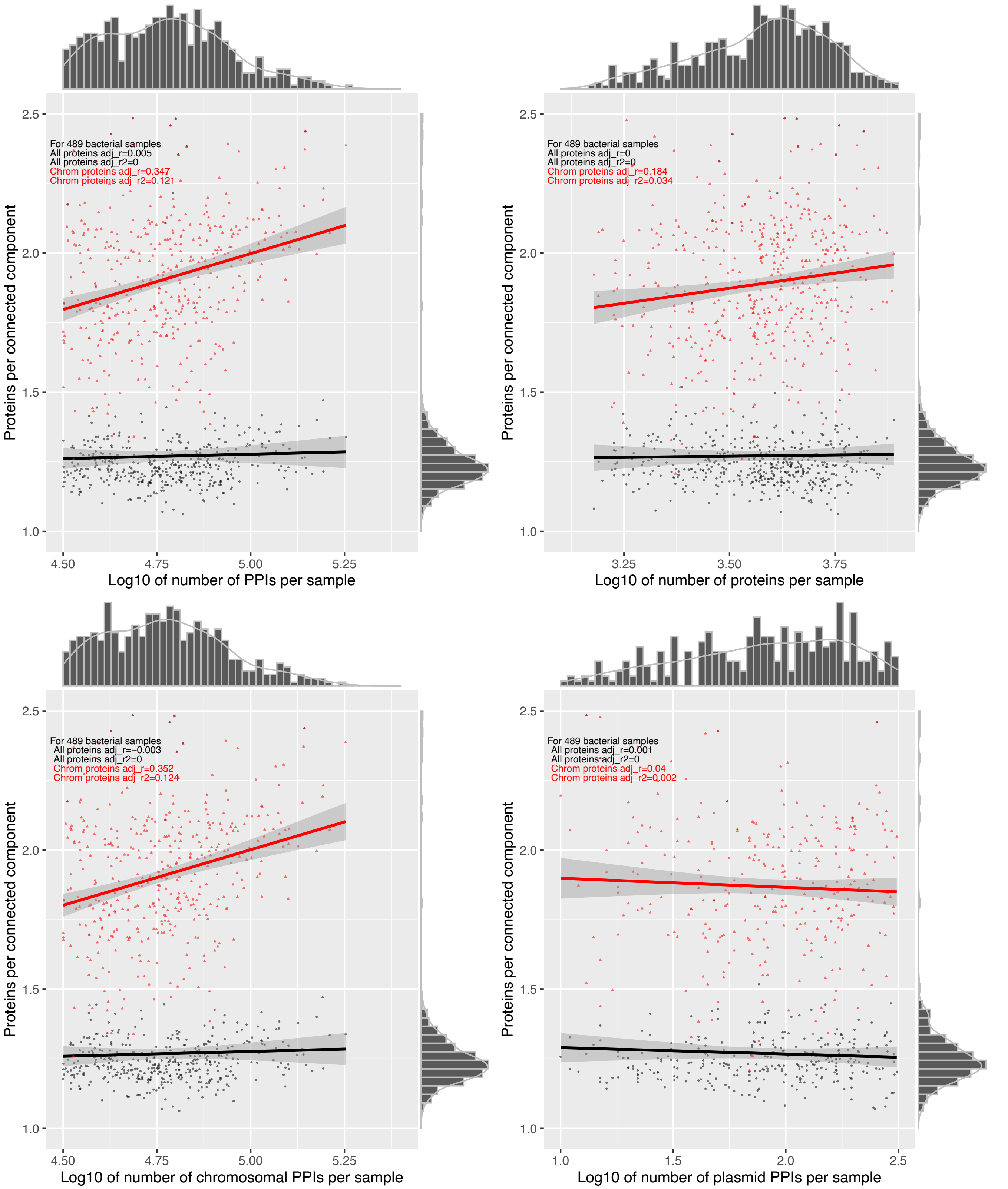


**Figure S23**. Varying levels of a positive correlation between the number of proteins per connected component (y-axis) for all proteins (black) and chromosomal proteins (red) with: (A) the log10-scaled number of PPIs, (B) log10-scaled number of plasmid-related PPIs, (C) log10-scaled number of chromosome-restricted PPIs, and (D) log10-scaled number of proteins per sample. The data plotted is for the 489 samples. The line of best fit (black for all proteins, red for chromosomal ones) shows a linear correlation of each variable with the number of indirect connections per protein. The histograms indicate the marginal densities per axis.
